# Live cell imaging of metabolic heterogeneity by quantitative fluorescent ATP indicator protein, QUEEN-37C

**DOI:** 10.1101/2021.10.08.463131

**Authors:** Hideyuki Yaginuma, Yasushi Okada

**Affiliations:** Laboratory for Cell Polarity Regulation, RIKEN Center for Biosystems Dynamics Research, RIKEN, Suita, Osaka, 565-0874, Japan; Department of Cell Biology, Department of Physics, the Universal Biology Institute (UBI), International Research Center for Neurointellience (WPI-IRCN), the University of Tokyo, Hongo, Tokyo, 113-0033, Japan

**Author notes:** Correspondence should be addressed to Yasushi Okada. Department of Applied Chemistry, The University of Tokyo, Hongo, Tokyo, 113-0033, Japan (Hideyuki Yaginuma).

**Keywords:** ATP concentration, single-cell measurement, metabolic heterogeneity, fluorescent indicator protein, live cell imaging

## Abstract

Adenosine triphosphate (ATP) is often referred as the energy currency of the cell. Yet, non-invasive, real-time, and quantitative measurement of its concentration in living mammalian cells has been difficult. Here we report an improved fluorescent ATP indicator protein, QUEEN-37C, which is optimized for measuring ATP concentration in living mammalian cells. Absolute value of the ATP concentration can be estimated from the ratiometric fluorescence imaging, and its accuracy was verified by the luciferase assay. Since QUEEN-37C enables the single-cell measurement of ATP concentration, we can not only measure its mean but its distribution in the cell population, which revealed that the ATP concentration is tightly regulated in most cells. We also noted the positive correlations in the ATP concentration among adjacent cells in epithelial cell sheet and mouse embryonic stem cell colonies. Thus, QUEEN-37C would serve as a new tool for the investigation of the single cell heterogeneity of metabolic states.

## Introduction

Adenosine triphosphate (ATP) is often referred to as the energy currency of the cell. It is a high-energy molecule that stores the energy and is used for almost all cellular activities. Its synthesis or energy metabolism should be balanced to its consumption to keep the cells in the steady state. In addition to this housekeeping role, the energy metabolism of each cell would need to respond dynamically to the changes in the cellular states. In fact, many previous and recent findings suggest tight relations between the cellular energy metabolism and the cell state changes such as cellular differentiation, reprogramming or cancer (Shyh-Chang 2013, Wellen, 2009, Moussaieff 2015, Ito 2014, Takubo 2013). Cellular heterogeneity of metabolic states would play important roles in such cell state transitions (Tonn et al., Commun Biol 2019), but single-cell analyses of metabolic states have been difficult. Recent advancement in the metabolic analyses has enabled the real-time and non-invasive monitoring of oxygen consumption (OCR: oxygen consumption rate) and glycolysis activity (ECAR: extracellular acidification rate), but they can only measure the metabolic activity of a cell population but not at the single cell level. Hence, cellular heterogeneity in the metabolic activities has been difficult to examine directly.

Live cell fluorescence imaging is one of the most suitable tools for the measurement of cellular heterogeneity. Fluorescent indicators for the metabolic states are required for such measurement. The fluorescent indicators for ATP concentration (Rangaraju 2014; Wang 2014; Morciano 2017; Berg 2009; Tantama 2013; Arai 2018) would be good candidates. Some of them have difficulties in the application for the single-cell measurement such as lower sensitivity, smaller dynamic ratio, or difficulties in introducing into cells. But more critical limitations are the difficulties in quantitative estimation of the ATP concentration. This is a common limitation with various fluorescent protein (FP)-based indicators (Nasu et al., Nature Chem Biol 2021). For example, the most successful examples are genetically encoded calcium indicators. They are not usually used for the measurement of the calcium concentration, but for the monitoring of the electric activity of neurons, where only the rapid increase of the intracellular calcium concentration needs to be recorded.

Ratiometric measurement with Forster resonance energy transfer (FRET)-based indicator is generally considered as more appropriate for quantitative measurement, and FRET-based ATP indicators have been previously developed to measure concentration of ATP at single cell level (Imamura 2009, Nakano 2011). These indicators enabled us to visualize the relative temporal or spatial changes within in a same cell (Ando 2012; Tanaka 2014; De Bock 2013; Pathak 2015). However, we noted that FRET-based indicators have intrinsic biases for the quantitative estimation of the ATP concentration (Yaginuma 2014). The apparent FRET efficiency can be affected by the differences in the maturation and degradation kinetics of FPs that constitutes the FRET pair. One of the two FPs in the pair might be missing due so the slower maturation or faster degradation. The linker between the FRET pair might be degraded leaving two separate FPs. The degree of these biases can be variable between cells, and careful calibration would be required for each cell, which would limit applications.

To circumvent this problem, we previously developed an ATP indicator named QUEEN (for QUantitative Evaluator of cellular ENergy). It is a single FP-based, ratiometric fluorescent ATP indicator optimized for the quantitative measurement of ATP inside bacterial cells at 25°C, which revealed unexpectedly large heterogeneity in the intracellular ATP concentration among the clonal bacterial cells in the same culture (Yaginuma 2014).

Here we report a new version of QUEEN optimized for the measurement in mammalian cells at 37°C. The concentration of intracellular ATP can be estimated from the pre-determined calibration curve. Its accuracy was validated by luciferase assay. As a demonstration for the application in the metabolic heterogeneity measurement, we measured the distribution of intracellular ATP concentrations in more than 20 commonly used cell lines or primary cells. To our surprise, most of the cell lines showed much smaller diversities in the ATP concentration, suggesting much tighter regulation than in bacterial cells. As an example for such regulatory systems, we also noted the spatial correlation of ATP concentration among adjacent cells in metabolically challenged epithelial cell sheet or in undifferentiated mouse embryonic cell colonies.

## Results

### Development of QUEEN variants

The intracellular ATP concentration normally ranges in 1-10 mM, and our previous indicator QUEEN was tuned to respond in this range at 25°C. The affinity of the sensor region was rationally engineered based on our previous studies for the development of ATP indicators based on the epsilon subunit of F_o_F_1_-ATP synthase (Kato-Yamada 2003, Yagi 2005, Imamura 2009, Yaginuma 2014). The epsilon subunit from *Bacillus* PS3 has higher affinity to ATP than that from *Bacillus subtilis*. The indicator with both N-terminal and C-terminal half of *Bacillus* PS3 (QUEEN-7μ) showed much higher affinity to ATP than physiological level (*K*_d_∼7 μM, Yaginuma 2014). Exchanging the N-terminal half (amino acids 1-107) with *Bacillus subtilis* lowered the affinity (QUEEN-2m, Yaginuma 2014) and responded well for the physiological ranges (1-10 mM) of ATP at 25°C.

Since the affinity of QUEEN-2m is too low for the measurement in the mammalian cells at 37°C while it is too high for measurement at 20°C, we made a series of QUEEN variants. We first created another chimeric indicator, QUEEN BP84 (amino acids 1-83 of *Bacillus subtilis* and amino acids 84-133 of PS3), which showed slightly higher affinity than the physiological range (*K*_d_∼1 mM). Next, we introduced mutations to QUEEN-7μ, QUEEN-2m and BP84 to change their affinity. The mutants were expressed in bacterial cells and examined the affinity to ATP in vitro (Supplementary Table S1). Thus we obtained two new QUEEN variants suitable for the measurement at 37°C (QUEEN-37C) and 20 °C (QUEEN-20C), respectively (Fig. 1A).

**Figure 1.**
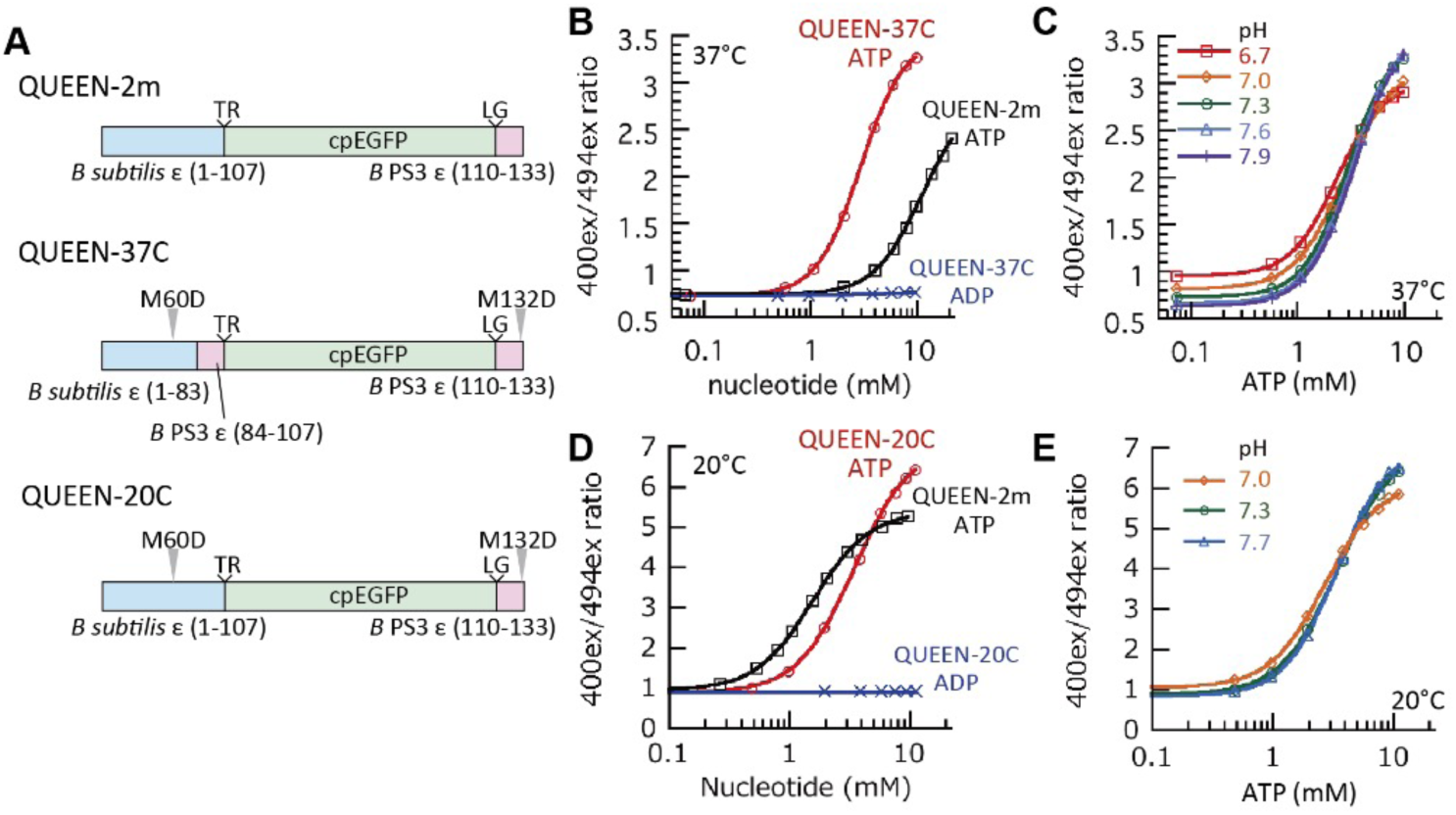
Development of QUEEN-37C. (A) Structures of previous and newly developed QUEENs. (B) Response curve of QUEEN-37C to nucleotides. (C) Sensitivity of QUEEN-37C response curve to pH. (D) Response curve of QUEEN-20C to nucleotides. (E) Sensitivity of QUEEN-20C response curve to pH.

Since these new variants have mutations in the sensor domain, we have re-evaluated the selectivity to ATP over adenosine diphosphate (ADP) and the sensitivity to pH changes. As shown in Figs 1 B-E, they both showed good selectivity to ATP over ADP, and little dependence to pH changes. The binding and dissociation rate constant of QUEEN-37C to ATP (k_on_ = 3.5 × 10^-2^ s^-1^ mM^-1^, k_off_ = 1.7 × 10^-1^ s^-1^) was similar to that of previous QUEEN-2m (Yaginuma et al, 2014). In the following sections, we focus on QUEEN-37C and its applications in mammalian cells. Further characterization of QUEEN-20C is beyond the scope of this report, but we expect that it would be suitable for the measurement around 20°C, for example with *C elegans*.

### Validation of the accuracy of ATP concentration measurement by QUEEN

Calibration is important for the quantitative measurement, but it is often difficult for the assays with living cells. If the response in the living cell is same as in vitro, the calibration curve pre-determined in vitro can be used for the live cell assays. To validate this idea, we compared the QUEEN-37C signals with the intracellular ATP concentration measured by luciferase assay (Fig 2).

**Figure 2.**
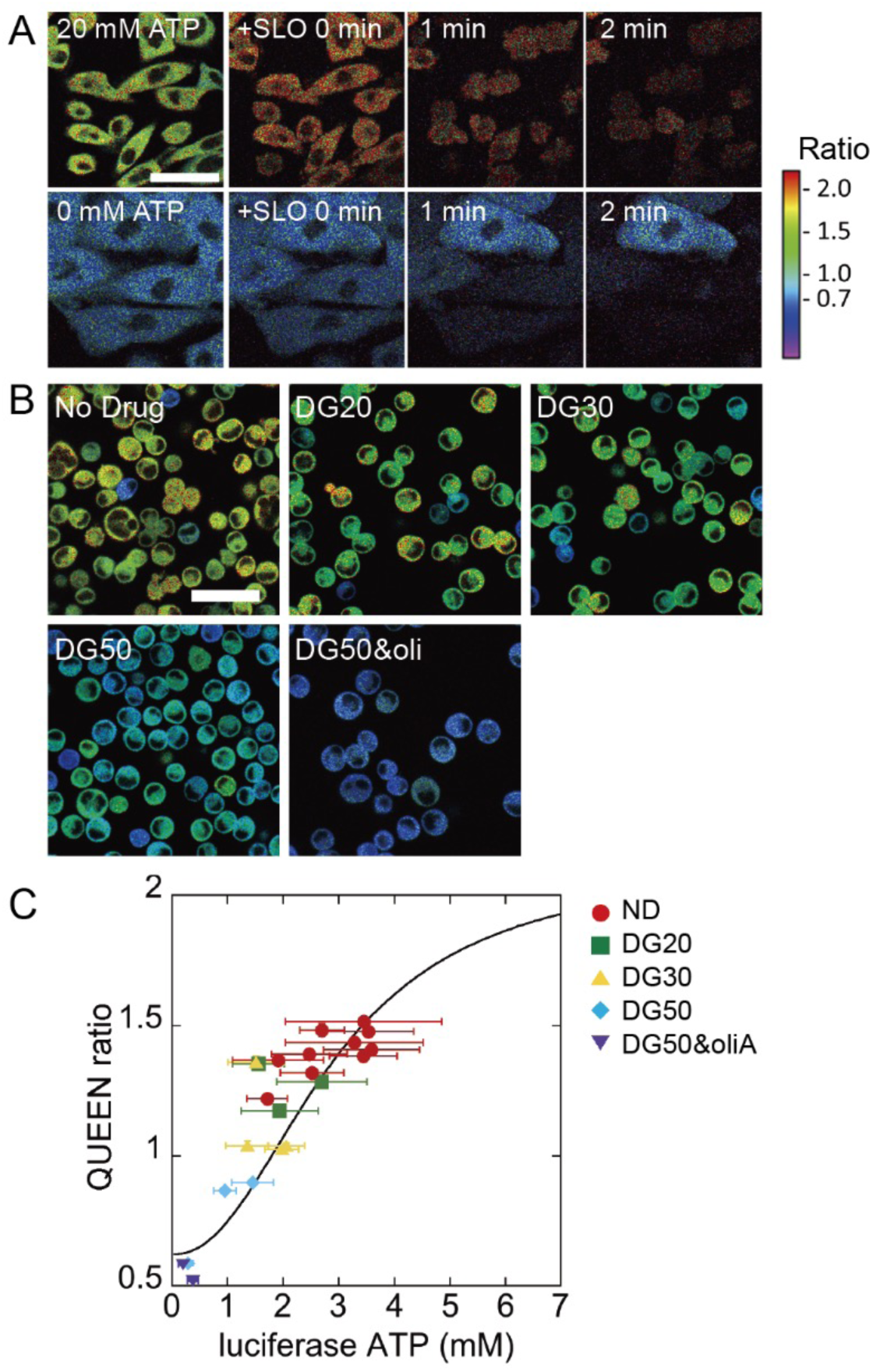
Measurement of the same samples with QUEEN-37C and luciferase assay. (A) Ratio image of cells during SLO treatment. Extracellular ATP concentration is 20 mM (top row) or 0 mM (bottom row). Scale bar = 50 µm. (B) Representative images of detached QUEEN-37C MDCK cells with or without various concentrations of 2-DG and oliA. Scale bar = 50 µm. (G) QUEEN-37C ratio plotted against cytosol ATP. Solid line is the theoretical relationship derived from in vitro measurements and SLO experiment in (A). ND: No drug, DG20: 20 mM 2-DG, DG30: 30 mM DG, DG50: 50 mM 2-DG, DG50&oliA: 50 mM 2-DG and 10 µM oliA.

We first calibrated the differences in the spectral properties of the instruments we used for the measurements in vitro (spectrofluorometer) and with living cells (confocal microscope), respectively. Purified QUEEN-37C protein was measured in vitro for the ratio of fluorescence at 400 nm excitation divided by 494 nm excitation (400ex/494ex) with spectrofluorometer. The results were fitted with Hill equation to estimate the minimum and maximum ratio values, *K*_d_, and Hill coefficient. Among these parameters, the former two values are dependent on the spectral properties of the instrument. The latter values reflect the biochemical properties of the sensor domain, which would not be affected by the instruments or measuring methods.

Next, we determined the maximum and minimum ratio values for the microscope observation by permeabilizing the plasma membrane of QUEEN-37C expressing cells with streptolysin O (SLO) with oversaturating concentration of ATP (20 mM) in the medium or without ATP (0 mM) as shown in Fig 2A. The ratio value jumped up or down immediately after permeabilization before the signal loss by diffusion of QUEEN protein from the cell. These values were used as the maximum and minimum ratio values. The calibration curve can be calculated from these values if we assume the *K*_d_ and Hill coefficient values do not change much in living cells than in vitro.

The validity of this estimated calibration curve was examined by comparing the microscope images of the QUEEN-expressing cells with the intracellular ATP concentrations measured by the luciferase assay after imaging (Fig 2B, C). The intracellular ATP concentrations were gradually lowered by partial inhibition of ATP synthesis with various concentrations of 2-deoxyglucose (2-DG). In some experiments, oligomycin A (oliA) was also added for further inhibition. The cell suspension was imaged with confocal microscope (Fig 2B), and then measured the population average of the ATP concentration inside the cells by luciferase assay. The determination of the cell volume, cell density, as well as the corrections for the systematic errors (for example, degradation of ATP during inactivation and solubilization of the cell) are detailed in the method section.

The results are plotted in Fig 2C. The points with error bars are the results of the measurements. They agreed well with the estimated calibration curve based on the biochemical parameters (*K*_d_ and Hill-coefficient) determined with purified protein in vitro. Namely, we can use this calibration curve to estimate the cellular ATP concentration from the images with QUEEN. It should be noted here that frequent calibration is not required for the routine measurement. Calibration is required only for the spectral response parameters (the maximum and minimum ratios), but they are dependent on the instrument and are stable over time.

The validity of this calibration method was thus confirmed experimentally. But there remains a possibility that acute inhibition of ATP synthesis might have caused some unphysiological changes in the cell. For example, ATP is proposes as a hydrotrope in the cell to prevent phase separation or protein aggregation in the cytoplasm (Patel et al, 2017). Its acute decrease might change the physicochemical properties of the cytoplasm to induce phase separation or aggregate formation. We examined this possibility by monitoring stress granule (SG) formation, a membrane-less organelle linked to protein aggregation. A SG marker protein G3BP1-iRFP was expressed along with QUEEN-37C to examine the SG formation and ATP concentration changes simultaneously. As shown in Supplementary Fig. S1, the formation of the phase separation granule occurs only in small population of cells and much later than the decrease of ATP concentration. Thus, the acute decrease of ATP would not cause large and immediate changes in the physicochemical environment of the cytoplasm. However, some ATP depleted cells formed SGs later, which might be consistent with the proposed roles of ATP in the regulation of phase separation or prevention of protein aggregation.

### Measurement of cellular heterogeneity of ATP concentration in various cell types

The average concentration of ATP in mammalian cells is reported to be around 3 mM (Traut, 1994). The exact value would be different among various cell types or the cellular states, but only average values have been examined in most studies to date. One might assume that ATP concentration would be tightly regulated because it is an essential energy currency, but our previous work with bacterial cells demonstrated even the clonal *E. coli* cells in continuously growing condition showed unexpectedly large variance (Yaginuma 2014). The SD value (1.22 mM) was comparable to the mean value (1.54 mM). Namely, the coefficient of variation (CV) value was 0.8. The cellular heterogeneity was much larger than expected in bacterial cells.

It is yet unclear whether mammalian cells show such large heterogeneity. Heterogeneity might be caused by the limited number of metabolic enzyme molecules in the small bacterial cells. We have therefore measured the heterogeneity of ATP concentration in various mammalian cells. More than 20 commonly used cell lines were obtained from the cell bank and QUEEN-37C was introduced for measurement (Fig. 3A, B, Table 1, Supplementary Dataset S2). The ATP concentration distributed mostly in the range of 3-7 mM. Cells with 2 mM or lower ATP were rare. Namely, the ATP level would be maintained above 2 mM in healthy mammalian cells. This makes a big contrast with our previous report in *E. coli* cells, where most cells were below 2 mM (Yaginuma 2014). The average ATP concentration was variable among cell lines; about 4 mM for many cells, but 6 mM or more for some cells (Table 1). More interestingly, most cell lines showed much smaller heterogeneity (CV 0.1∼0.2) than bacterial cells. Some cells showed larger heterogeneity, for example in primary cultured neuron (CV∼0.55). This might reflect the heterogenous mixture of cell types in the primary culture. These results show that QUEEN-37C enables the simple and easy measurement of metabolic heterogeneity by imaging. Such measurement would enable future studies to clarify the origin and physiological relevance of the metabolic heterogeneity.

**Figure 3.**
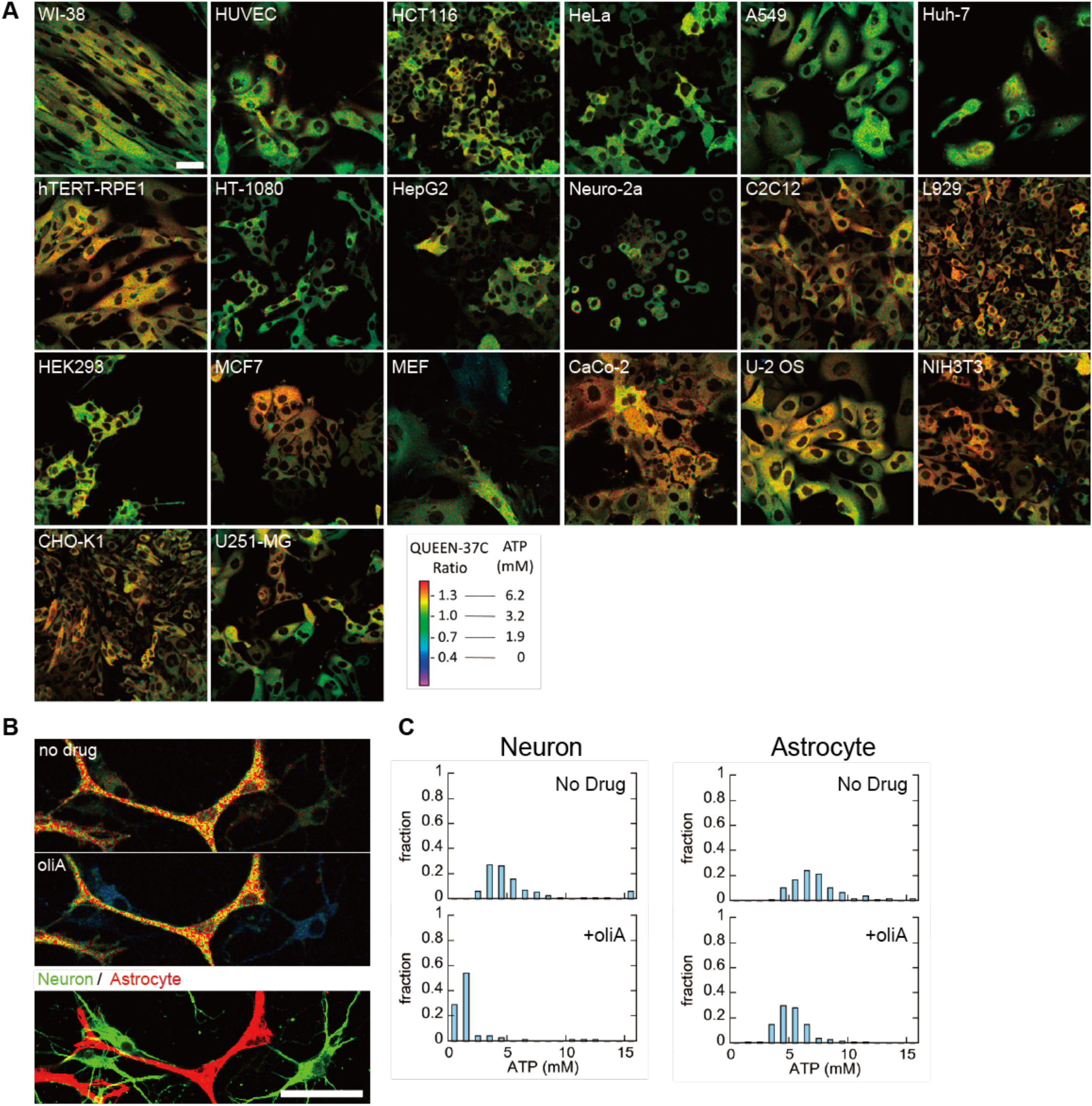
Meaurement of various cell lines and primary cells. (A) Representative ratio images of measured cells, except for neuron and astrocyte. Scale bar = 50 µm. (B) Response of mouse hippocampal primary neurons and astrocytes to oliA. Ratio images before (top left) and after (bottom left) oliA treatment are shown. Immunostaining image to discriminate between neurons and astrocytes are shown on the right. Scale bar = 50 µm. (C) Histograms of measured ATP values in neurons (left) and astrocytes (right).

**Table 1.**
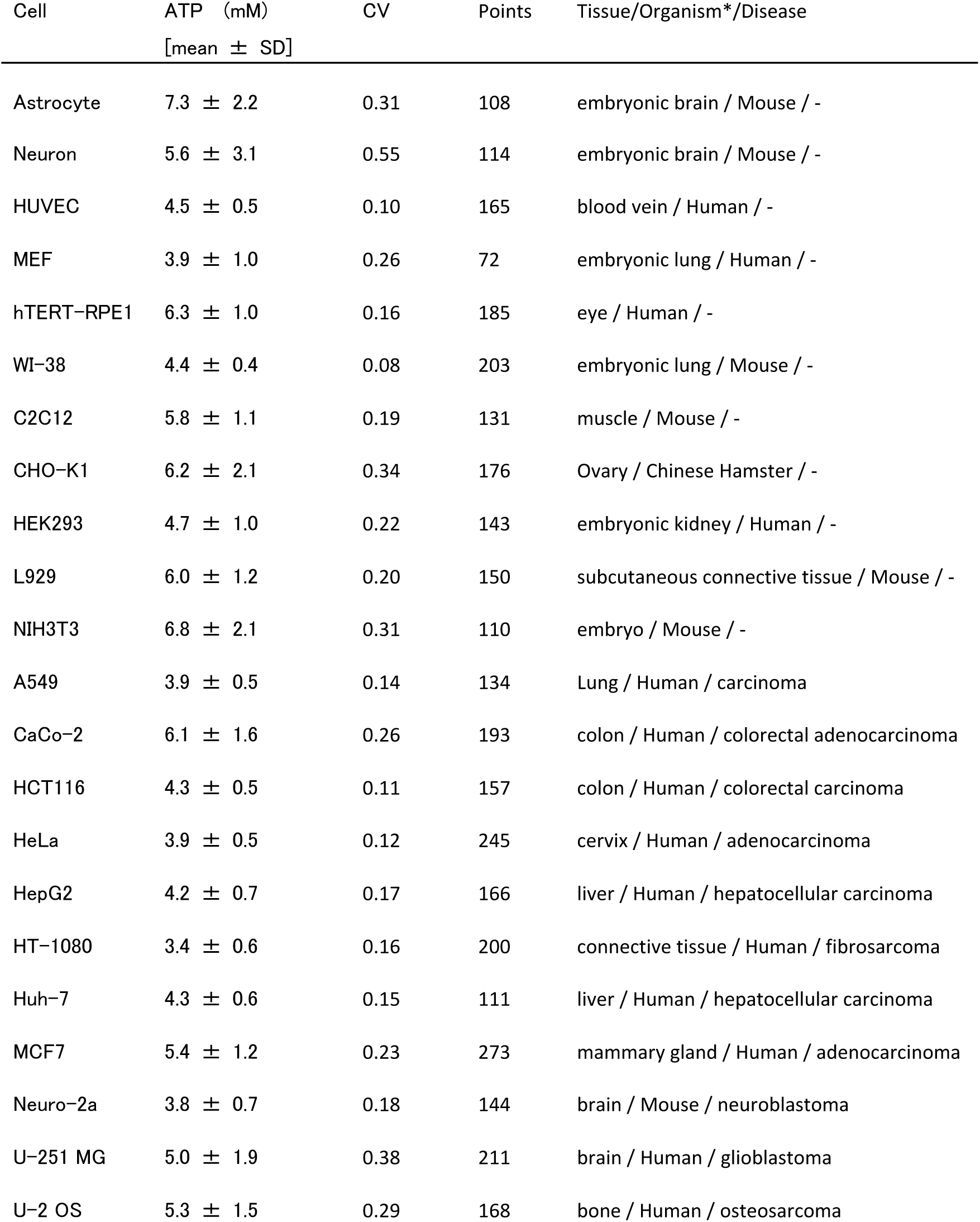
Summary of the distribution of single-cell ATP levels in various cell types measured using QUEEN-37C.

### Dynamic response to metabolic perturbations

The merit of ATP measurement with live imaging is not limited to the measurement of the cellular heterogeneity at a specific time point. Temporal responses to perturbations would enable us to infer the underlying metabolic processes. As a proof of its concept, we examined the responses to the inhibitions to aerobic glycolysis (hereafter we refer as glycolysis for simplicity) or oxidative phosphorylation (OxPhos), the two major pathways for cellular ATP synthesis. The cells mainly dependent on glycolysis would respond sensitively to its inhibition, and vice versa. Thus, we can infer whether or how much the target cells are dependent on glycolysis and OxPhos.

We first examined the mixed primary culture of neurons and astrocytes, because neurons are mainly dependent on OxPhos, while astrocytes are mainly glycolytic (Belanger, 2011). Application of oliA, an inhibitor of the OxPhos enzyme F_o_F_1_-ATPase, only affected slightly to astrocytes, but neurons decreased ATP drastically (Fig. 3 B and C). These results would confirm that the metabolic states or the balance between the glycolysis and the OxPhos can be examined at a single cell level by using the combination of QUEEN-37C and the inhibitors.

Similar results were obtained with a canine kidney cell line, MDCK (Madin-Darby Canine Kidney) cells (Fig 4). Some cells responded sensitively to oliA, while others are more sensitive to oxamate, an inhibitor for the glycolytic enzyme lactate dehydrogenase.

**Figure 4.**
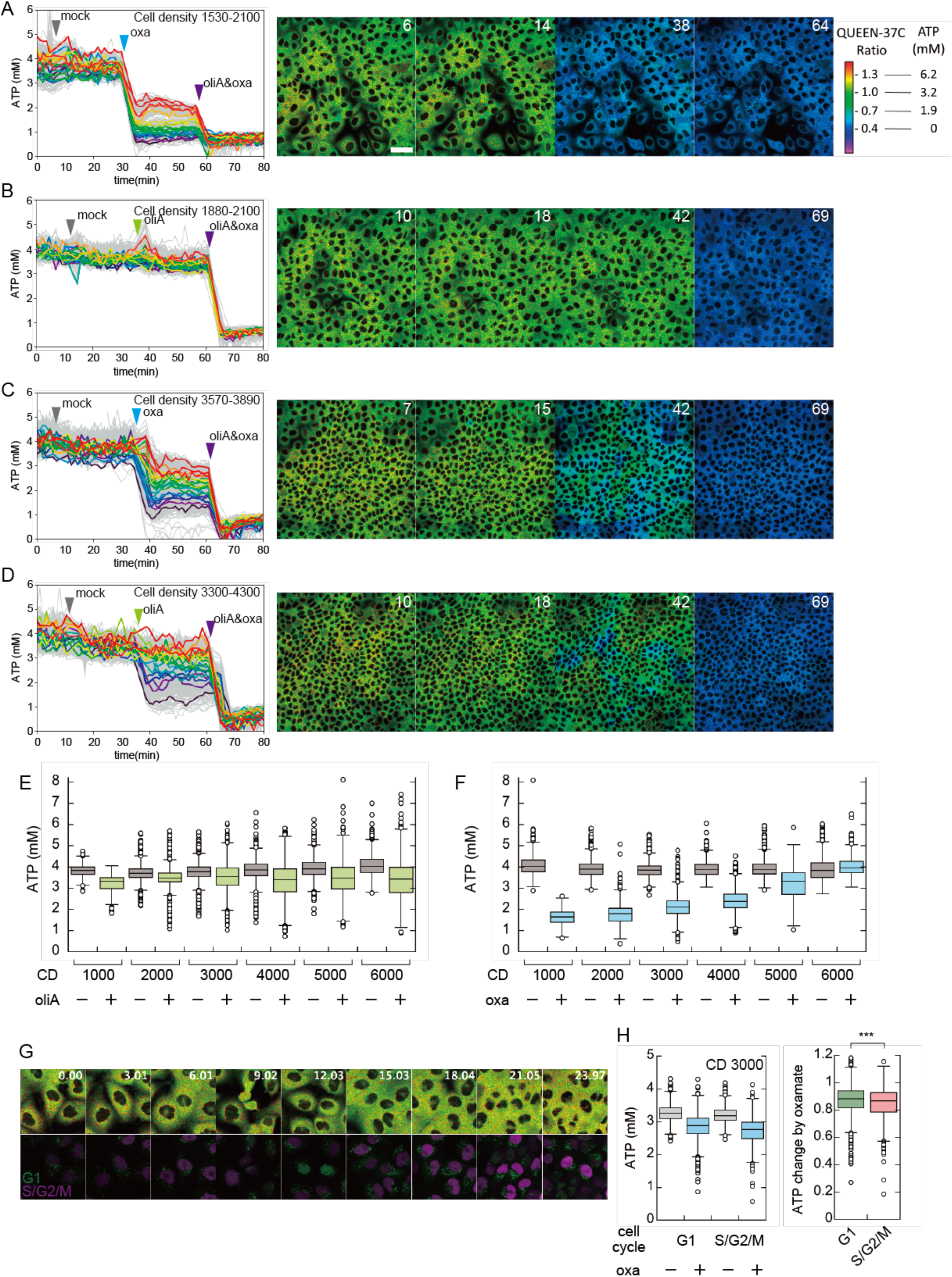
Response of MDCK cells expressing QUEEN-37C to energy metabolism inhibitors. (A-D) 1 min-interval timelapse measurement of MDCK cells during ATP synthesis inhibitor treatment. Cells were first treated with mock medium, then with either oliA (A, C) or oxamate (B, D), and finally with both drugs. Low cell density region (A, B) and high cell density region (C, D) were plotted separately. Time course is on the left side, and representative images are shown on the right. In the time course plot, each single line represents individual cells. For clarity, 20 plots were selected and highlighted in different colors, and other plots are shown in gray. To select the colored plots, cells were sorted by ATP concentration after first drug treatment, and 20 cells were selected at regular intervals. Scale bar = 50 µm. (E-F) Response to oliA (E) or oxamate (F) treatment at various cell density. CD = cell density. (G) Images from Simultaneous imaging of QUEEN-37C and FUCCI cell cycle reporters, namely mScarletI-Cdt1 and iRFP-Gem. (H) Oxamate sensitivity of MDCK cells in G1 phase and S/G2/M phase. Box whisker plots of absolute ATP concentrations before and after oxamate treatment (left), and ATP concentration drop ratio by oxamate treatment (right).

The observed heterogeneity in the responses might reflect the metabolic heterogeneity of the cells, namely whether the cell is more dependent on glycolysis or OxPhos. However, QUEEN signal might be affected by the inhibitor drugs. In fact, the ATP response of QUEEN changes at pH conditions below 7.0 (Fig 1C), which can occur after metabolic inhibition (Tantama et al, 2013, Supplementary Fig S2 A-C).

We, therefore, measured ATP and pH simultaneously in MDCK cells. Ratiometric pH indicator protein pHluorin (Miesenbock et al 1998) was expressed as a fusion protein with nuclear localization signal so that the pH signal is separated from the cytoplasmic QUEEN signal. In a separate experiment, we confirmed nuclear pHluorin showed the same response to cytoplasmic pHluorin (Supplementary Fig S2C). Only cells expressing high level of nuclear pHluorin were measured for unambiguous separation of the two signals (Supplementary Fig S2D). Even in the highly expressing cells, the leakage of nuclear pHluorin to the cytoplasm was relatively small (Supplementary Fig S2E). In order to minimize the effect of leakage, QUEEN-37C ratio was calculated after subtracting the estimated leakage fluorescence. We can thus measure pH and QUEEN signal simultaneously in the same single cells. As shown in Supplementary Fig S2 F-H, the intracellular pH showed slight decrease after oxamate or oliA application. The pH drop was larger with lower density cultures, but still the intracellular pH was above 7.0 (Supplementary Fig S2 F, G), and its effect on QUEEN is almost negligible.

It should be noted however, that application of both oliA and oxamate lowered pH to about 6.7. In this acidic environment, QUEEN does not respond sensitively to the decrease of ATP (Fig 1C). The ATP concentration will be estimated to be higher than the real value in this condition.

Simultaneous measurement of ATP and pH was difficult in ATP depleted cells, because the nuclear localization of pHluorin is no longer maintained. We, therefore, report the estimated ATP concentration without pH correction. The QUEEN signal (405ex/488ex ratio) did not drop enough after the double inhibition with oliA and oxamate, but the true ATP concentration would have decreased to nearly 0 mM.

### Cell density dependent metabolic switching in MDCK cells

Through the above experiments, we noticed that MDCK cells showed heterogenous responses to oliA or oxamate. MDCK cell is a widely used model system for epithelial cell polarity development (Dukes et al, 2011). When cultured at lower densities, MDCK cells take a flat and unpolarized shape. At higher densities, however, they take a tall and cylindrical shape with clear apical-basal polarity and form E-cadherin dependent adherence junction. These features are typical morphological differentiations into an epithelial cell monolayer (Balcarova-Standar, 1984). We surmised that the major source of ATP synthesis in MDCK cells would also change from glycolysis to OxPhos during the epithelial development.

For this purpose, we have established a clonal MDCK cell line stably expressing QUEEN-37C, and measured the time-dependent changes of the signal after the application of oliA or oxamate. The cells were first measured without inhibitors, then either of the two inhibitors were applied to measure the response. Finally, both inhibitors were added to block both pathways completely.

As expected, low density culture (∼2000 cells/mm^2^) responded sensitively to oxamate but very little to oliA (Fig 4A, B). Almost all cells reduced ATP concentration more than 50% by the application of oxamate, but very few cells showed decrease by oliA. Contrastingly, higher density culture (∼4000 cells/mm^2^) showed more heterogeneous responses (Fig 4C, D). Some cells showed resistance to oxamate, while many other cells responded (Fig 4C). Consistently, some cells responded to oliA, while many others were still resistant (Fig 4D). The population of the oxamate-responding cells or oliA-resistant cells appeared to increase at even higher densities (Fig 4 E, F, Movie S2, S3, Dataset S3). We examined if the apparent cell size in the image is correlated with the response to oxamate or oliA. We found that smaller cells showed significantly more response to oliA and less response to oxamate as a whole. However, when we took a closer look, smaller cells showed a variety of response to both metabolic inhibitors (Supplementary Fig. S3). This suggests that response to oxamate and oliA is not dictated by apparent cell area.

The ATP concentration dropped below 1 mM after application of both oliA and oxamate (Supplementary Fig. S4). The true ATP concentration would be much lower as discussed above. This would suggest that OxPhos and glycolysis are the two major sources for ATP synthesis in these cells and switched the dependency from glycolysis to OxPhos during the epithelial differentiation. Here we noted that the final ATP concentration tended to be higher in higher density cultures (>4500 cells/mm^2^), which might suggest the presence of the third ATP synthesis pathway.

### Metabolic heterogeneity in MDCK cells

MDCK cells showed heterogenous response to oliA or oxamate at nearly confluent cell densities (∼4000 cells/mm^2^). Cell cycle would be one possible source for this heterogeneity. We tested this idea by establishing a MDCK cell line that stably expresses QUEEN-37C and FUCCI-cell cycle markers: a S/G2/M phase marker, miRFP709-hGem and G1 marker, mScarletI-hCdt1 (Shchbervacova et al, 2016). The FUCCI-cell cycle markers worked correctly in the QUEEN-37C cells (Fig. 4G), and the response to oxamate was evaluated for mScarletI-hCdt1 positive G1 cells and negative S/G2/M cells. G1 cells were slightly less sensitive to oxamate (Fig 4H), but the difference was much smaller than the observed heterogeneity. The cell cycle itself would, thus, have relatively smaller effects, if any, on oxamate sensitivity.

During these inhibitor experiments with MDCK cells, we noticed a tendency that neighboring cells seemed to show similar ATP responses. OliA-responding cells often make clusters or islands in non-responder cell sheets (Fig 4C). Oxamate-resistant cells also make similar islands (Fig 4D).

We tested this tendency in a quantitative manner by autocorrelation analysis. Images after oliA or oxamate inhibition at ∼4000 cells/mm^2^ were analyzed (Fig 5). Coarse-grained ATP concentration images were created by 8×8 median binning, and radially averaged autocorrelation function was calculated. Two shuffle images were produced as a control (Fig 5 C, D, G, H). The pixel values are shuffled within each cell regions (“shuffle inside cell”) or the pixels in each cell region is exchanged with the pixels from another randomly selected cell region (“shuffle cells”). The latter shuffling will erase the spatial correlations between the adjacent cells, but the former shuffling will not. In fact, the correlation dropped sharply at around 10 µm with “shuffle cells” images, while “no shuffle” or “shuffle within cells” images showed correlation over much longer distance about 50 µm (Fig 5 I, J). The 10-µm correlation in the “shuffle cells” images would reflect the size of each cell. The correlation distance of 50 µm corresponds to about 2-3 cells, which quantitatively supports the apparent cell islands with about 5 cells in diameter. These islands might be produced from the daughter cells of progenitor cells. Alternatively, epithelial cells are connected physically and chemically. Signaling at cell-cell junctions might have synchronized the responses between the adjacent cells.

**Figure 5.**
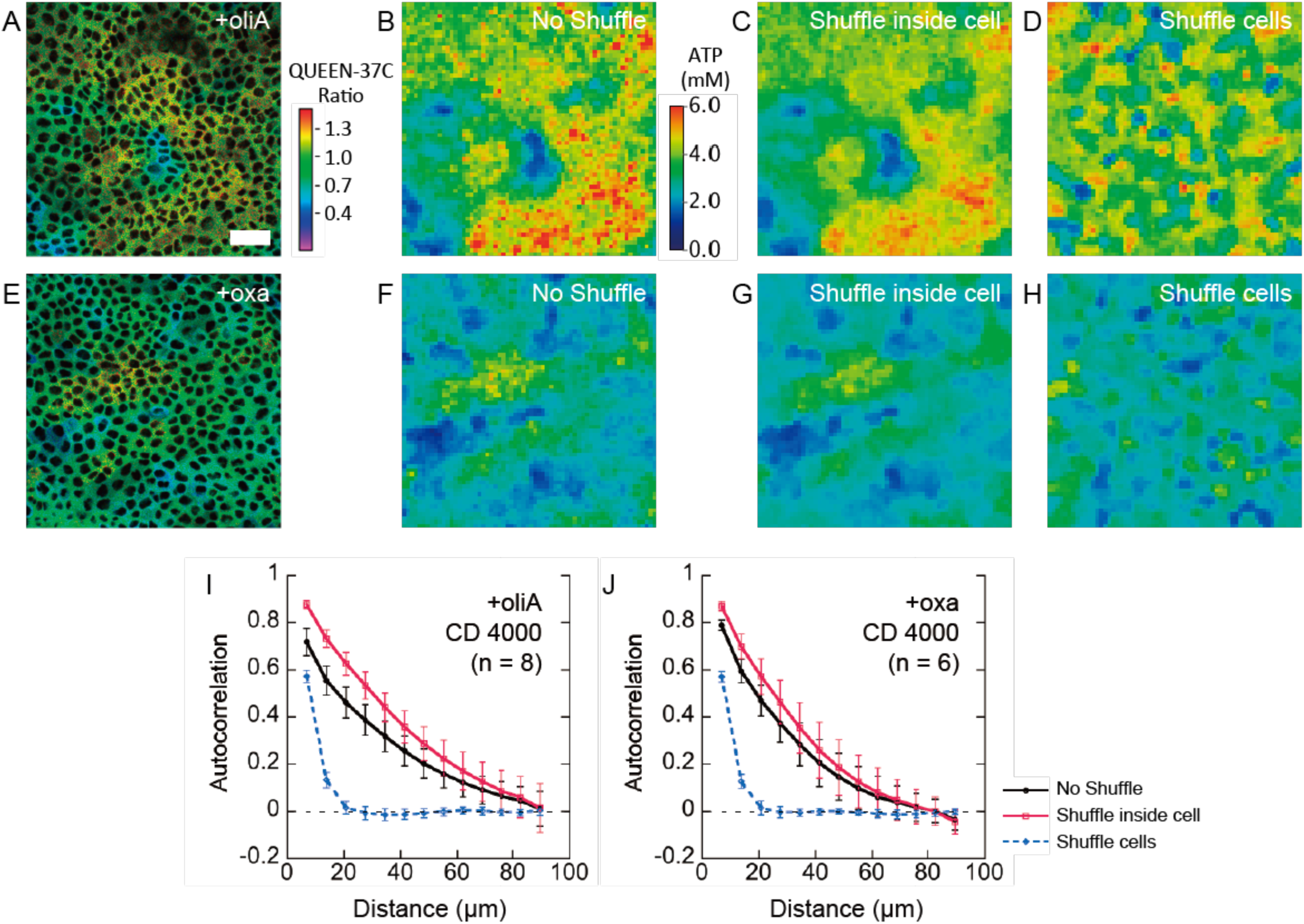
Spatial autocorrelation analysis of ATP values after drug treatment. (A, E) QUEEN ratio image obtained from experiment. Brightness is scaled to fluorescence intensity. Scale bar = 50 µm. (B, F) Coarse-grained image of ATP value distribution created from experimental images (A) and (E). (C, G) Computer-simulated image of ATP concentration created from (A) and (E). Ratio values in each pixel were shuffled within the same cell and then coarse-grained ATP concentration image was created. (D, H) In addition to pixel shuffling in each cell, the pairing of cell position and ratio values was also randomized. (I, J) Radially averaged autocorrelation of ATP values in experimental and simulated ATP concentration images. Several image fields (the number is descripted in the plot) were analyzed in the same manner, and the average autocorrelation at the corresponding distance is plotted. Bar = SD. (A-D, I) MDCK cells treated with oliA. (E-H, J) MDCK cells treated with oxamate.

### Metabolic heterogeneity of mouse ES cells

As another model of cellular heterogeneity, we examined mouse embryonic stem (mES) cells. We introduced QUEEN-37C to mES cells and maintained in the medium containing serum and leukemia inhibitory factor (LIF). In this condition, mES cells are known to show large heterogeneity in the expression level of the pluripotency-associated genes (Toyooka et al 2008; Wray et al 2011). As expected from this large heterogeneity, the ATP level of the QUEEN-mES cells showed large diversities ranging from 2.0 mM to 5.0 mM (Fig. 6A). More interestingly, the diversity between the mES cell colonies was significantly larger than within the colonies (p < 0.001).

**Figure 6.**
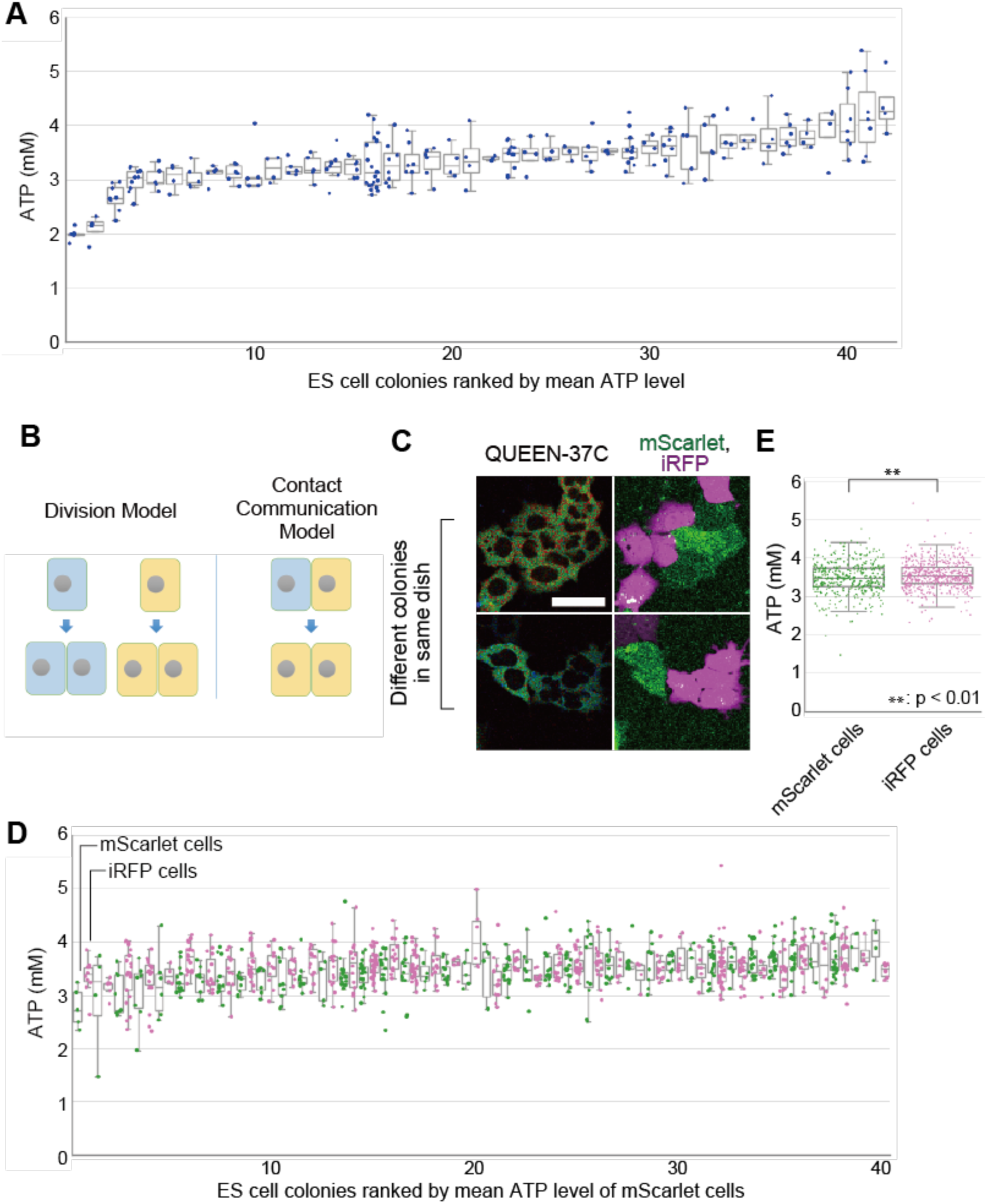
ATP measurement in mES cell colonies. (A) Single-cell ATP levels distribution in mES cell colonies. Colonies are shown in the order of mean ATP values. (B) A cartoon scheme for the two models that explains similar metabolic states between adjacent cells. (C) QUEEN-37C ratio image (left) and mScarlet/iRFP overlay image (right) of two representative colonies in the co-culture of two cell lines. (D) Single-cell ATP levels distribution in mES cell colonies in the co-culture of mScarlet and iRFP cell lines. Colonies are shown in the order of mean ATP values of the mScarlet cell lines. (E) Single-cell ATP levels of two cell lines, aggregated from 40 colonies in (D). The difference between two cell lines was significant (p < 0.01, Student’s t-test) but very small.

These observations might imply that mES cells in the same colony might communicate with each other and synchronize their differentiation states and thus their ATP levels might be synchronized (“communication model”). Alternatively, cells might show similar metabolic states simply because these cells are daughters from a common ancestor (“progenitor model”) (Fig. 6B). To distinguish these possibilities, we labeled QUEEN-expressing mES cells with two colors, mScarlet and iRFP. We mixed these cells and examined the colony of a mixture of two colors (Fig. 6C). Apparently, ATP level of iRFP cells was correlated with that of mScarlet cells in each colony (Fig. 6D). By analyzing single-cell ATP level with two-way ANOVA (Table 2), we confirmed that most of the variation in single cell ATP concentration can be explained by difference between colonies (p < 0.001). In other words, variation inside the colony is relatively small, even when different cell lines are forming a colony together. Therefore, this result is consistent with the communication model. Different cell line was also significant source of variation in ATP concentration (p < 0.001), but the difference between mScarlet and iRFP cell lines was very small and therefore might be trivial (Fig. 6E). The interaction between colony and cell line was a significant source of single-cell ATP variation (p < 0.001), too. This may reflect the observation that in some colonies the two cell lines shows different ATP concentrations (Fig. 6D), and this part of the result is supportive of the progenitor model. We believe that the two models are not mutually exclusive, and that both mechanisms would be involved in determining the metabolic state of the cell. Overall, our results would suggest the existence of some mechanism(s) that coordinates the ATP concentration between mES cells.

**Table 2.** Two-way ANOVA results for single cell ATP concentration in the colonies of two mES cell line co-culture.

| Source of Variation | Df | Sum Sq | Mean Sq | F value | Prob>F |  |
| --- | --- | --- | --- | --- | --- | --- |
| colonies | 39 | 19.49 | 0.50 | 4.10 | 0.000 | *** |
| cell lines (mScarlet or iRFP) | 1 | 2.17 | 2.17 | 17.82 | 0.000 | *** |
| colonies * cell lines | 39 | 9.85 | 0.25 | 2.07 | 0.000 | *** |
| Residuals | 946 | 115.25 | 0.12 |  |  |  |

## Discussion

We have developed a new ATP indicator, QUEEN-37C. QUEEN-37C enables quantitative measurement of ATP concentration at 37°C, the temperature suitable for use in most mammalian cells. In theory, QUEEN can avoid the biased signal problems of FRET-type indicators resulting from maturation/degradation kinetics of two fluorescent proteins, because QUEEN contains only one fluorescent protein (Yaginuma et al, 2014). Experimentally, the accuracy of QUEEN was confirmed by comparing the intracellular signal with luciferase assay. As is common to single FP-based sensors (Baird 1999), QUEEN-37C signal is mildly sensitive to pH. However, we showed here that in physiological and relatively mild experimental conditions, intracellular pH was within the range of 7.0-8.0. The effect of pH change to the QUEEN-37C signal in this range is negligible. Still, when quantitative measurements under more harsh conditions are required, the effects by the pH change would need to be compensated, for example, by the simultaneous pH measurement as described in this paper.

One of the advantage of QUEEN over other ATP indicators is that it can determine the ATP concentration in single cells rather than the relative changes over time. As its demonstration, we measured the cellular distribution of ATP concentration in more than 20 different common cell types. The mean ATP concentration ranged within 3-7 mM. Previously, the ATP concentration inside mammalian tissues and cells was reported to be around 3 mM (Traut 1994). Similarly, ATP amount in mouse or rat brain sample was determined to be 9-18 nmol / mg protein (Pissarek 1999, Manfredi 2002), which corresponds to approximately 2-4 mM, assuming that protein weight per cell volume is 0.2 g/mL (Milo 2013). Our results are consistent with these previously reported values, though they are slightly higher. This small discrepancy might be attributed to differences in sample preparation. We measured living cells on the microscope, whereas in previous experiments, cells and tissues were processed before measurement. Such sample preparation steps might have compromised regular metabolism and have caused mild decrease of ATP concentration.

We also showed that QUEEN-37C measurement can be applied to the single-cell analysis of the balance between glycolysis and OXPHOS. Such metabolic balance has been examined by measuring oxygen consumption or extracellular acidification (Nicholls 2010; Zhang, 2012), but it has been difficult to examine at the single cell levels. In this paper, we first examined with the mixed primary culture of neuron and astrocyte. It is well known that neurons are more dependent on OXPHOS, while astrocytes are primarily dependent on glycolysis (Belanger 2011). Various indicators have been used to demonstrate this difference such as ATP:ADP indicator, glucose indicator, NAD(P)H autofluorescence or FRET-type ATP indicator (Tantama 2013; Bittner 2010, Thevenet 2016). Our results with QUEEN-37C were consistent with these previous results. Moreover, QUEEN-37C enables quantification. The OXPHOS inhibition reduced ATP concentration by about 80% in neurons and 30% in astrocytes. These results confirm that neurons rely heavily on OXPHOS. The results also imply that OXPHOS contributes in a non-negligible proportion in astrocytes.

The same analysis was applied to an epithelial cell model, MDCK cell to examine the cell density dependent changes. Rapidly proliferating cells, especially stem cells, are known to be relying mainly on glycolysis, while more mature, differentiated cells are often dependent on OXPHOS for ATP production (Kondoh et al 2006; Vaurm et al 2011;Shyh-Chang 2013a; Meleshina et al 2016; Flores et al 2017). In our results, MDCK cells are much more glycolytic at low density when they are less differentiated and rapidly proliferating. This is in line with the above previous reports. Several mechanisms are proposed for this phenomenon, such as the increased demand for glycolytic intermediates as biomass substrates in proliferating cells (Vander-Heiden et al 2009). Alternatively, the concentration of metabolic intermediates might be coupled with cell fate through transcriptional regulation of genes (Shyh-Chang 2013b; Wei et al 2018). At the highest cell density in our study, treatment with both OXPHOS inhibitor and glycolysis inhibitor did not completely deplete ATP. We suspect that the inhibitor combinations we used did not stop all the possible ATP synthesis pathways. For example, sodium oxamate normally stops the glycolytic synthesis of ATP by inhibiting the reaction of lactose dehydrogenase (LDH) and preventing NADH to be converted back to NAD^+^. However, it might be possible to convert NADH back to NAD^+^ without using LDH or mitochondrial ATP production, for example by proton leakage of the mitochondria. Rapid uncoupling of mitochondria from ATP production for ATP homeostasis is reported previously (Wang, elife, 2017).

More interestingly, we found that the response of MDCK cells to inhibitors was spatially correlated. We found similar spatial correlation in mES cells. These results suggest the presence of cell-cell communication to keep the metabolic states of adjacent cells similar, although the mechanism for this is currently unknown. Gap junctions would be a good candidate, but the MDCK cell strain used in this study (MDCK II) reportedly does not form gap junctions (Dukes, 2011). It is also reported that mES cells can form gap junctions (Worsdorfer 2008). Spatial correlation of ES cell states are also reported in gene expressions (Okamoto, 2018), which would suggest the presence of some communication mechanisms between adjacent mES cells. The molecular mechanisms for this communication would require future research, but they should be important for a multicellular organism to synchronize cell fate between neighboring cells during development.

In conclusion, our new fluorescent ATP indicator, QUEEN-37C, is suitable for single-cell quantification of ATP concentrations. We propose that single-cell quantitative ATP determination is a useful non-destructive tool to study energy regulation processes that are blurred under previous ensemble and qualitative observations.

## Acknowledgements

We would like to thank Hiromi Imamura (Kyoto University), Akira Takai (BDR, RIKEN), Tetsuro Ariyoshi (BDR, RIKEN) and Daisuke Ino (Kanazawa University) for providing genetic resources, technical assistance, and helpful discussions. iRFP plasmid was a gift from Dr. Michiyuki Matsuda. We also thank the generous gift of plasmids through depositing to Addgene: Dorus Gadella, (#85042 and #85044), Vladislav Verkhusha (#80006 and #80007) and Didier Trono (#12259 and #12260). We also would like to thank Junko Asada, Shang-Dan. Xu, Nozomi Furutani, Manaho Kakiuchi and Tomoko Furuya for their technical and secretarial assistance. This work was supported by the Ministry of Education, Culture, Sports, Science and Technology through a Grant-in-Aid for Scientific Research (KAKENHI grant 15K18522 and 18K14721 to H.Y.; 16H05119, 19H03394, 19H05794, and 19H05795 to Y.O.), JST (JPMJCR15G2, JPMJCR1852, JPMJCR20E2, and JPMJMS2025-14 to Y.O.), the Takeda Science Foundation (Y.O.), and RIKEN SPDR program (H.Y.).

## Materials and Methods

### Reagents

Potassium chloride, sodium chloride, sodium diphosphate, disodium phosphate, potassium hydroxide, imidazol, glycerol, magnesium sulfate, calcium chloride, glucose, dimethyl sulfoxide (DMSO), fluorescein, 2-Amino-2-hydroxymethyl-1,3-propanediol (Tris) was purchased from Wako Chemicals. ADP, Isopropyl *β*-D-1-thiogalactopyranoside (IPTG). streptolysin O, olygomycin A, sodium oxamate, polyethyleneimine, poly-L-lysine was purchased from Sigma-Aldrich. 2-[4-(2-Hydroxyethyl)-1-piperazinyl]ethanesulfonic acid (HEPES), Piperazine-1,4-bis(2-ethanesulfonic acid) (PIPES) and ethylenediaminetetraacetic acid (EDTA) was purchased from Dojindo. ATP, guanine triphosphate (GTP), Triton X-100 was purchased from Nacalai Tesque. Puromycin, pencillin, streptomycin, fetal bovine serum, horse serum, TrypLE Express, non-essential amino acids (NEAA) solution, pyruvate solution, Glutamax solution was purchased from Thermo Fisher.

ATP, ADP and GTP were dissolved in water at approximately 300 mM and the pH was adjusted to 7.5 by adding potassium hydroxide solution. The concentration was determined by absorbance (ε = 15,400 M^-1^ cm^-1^ at 260 nm for ATP and ADP. ε = 13,700 M^-1^ cm^-1^ at 252 nm for GTP). Then the solution was separated in aliquots and stored at -80°C.

### DNA constructs for bacterial expression

We deposited previous QUEEN plasmids and the plasmids we created in the course of study to Addgene together with sequence information (See Supplementary Table S2). Starting from previously developed QUEEN-7µ (Addgene #129311) and QUEEN-2m (Addgene #129350) sequences, we first created an intermediate QUEEN variant “BP84” (Addgene #129378) using In-Fusion HD Cloning Kit (Clontech). BP84 is a chimeric protein of 1-83 amino acid sequence of *Bacillus. Subtilis* F_o_F_1_ *ε*, 84-107 of *Bacillus*. PS3 F_o_F_1_ *ε*, circularly-permuted enhanced GFP and 110-133 of *Bacillus*. PS3 F_o_F_1_ *ε*. Purified BP84 protein showed ratiometric fluorescence change to ATP concentration, but its *K*_d_ was ∼1 mM (Supplementary Table S1) and its response to physiological ATP quantification at 37°C was relatively small. To adjust the affinity at 37°C, we introduced one or two mutations in QUEEN-7µ and BP84, and created a series of candidate QUEENs. We expressed these variants in *E. coli*, obtained purified proteins and evaluated their response to ATP. The results of these mutants are also shown in Supplementary Table S1. Among these variants, A modified variant of BP84 with 2 additional single amino acid mutations, namely Y60D and M132D, showed good response to physiological ATP concentration range at 37°C, and was named QUEEN-37C (Addgene #129312). To create QUEEN-20C (Addgene #129316), a variant of QUEEN for usage at 20°C, the same mutations Y60D and M132D were introduced into QUEEN-2m to reduce the affinity at 20°C.

### Protein purification

QUEEN variants were purified from *E. coli* culture as previously described (Yaginuma et al, 2014) with some modifications. Briefly, JM109 (DE3) *E. coli* cells were transformed with pRSET B or equivalent vectors with QUEEN sequence. Cells were first cultured to OD 0.2-0.3 at 37°C, induced expression of QUEEN by 40 μM IPTG and cultured overnight at 18-28°C. Cells expressing QUEEN was pelleted and then suspended in the purification buffer [100 mM NaP_i_ (pH 8.0), 200 mM sodium chloride, 10 mM imidazol]. This cell suspension was sonicated, centrifuged and supernatant was collected. His-tagged QUEEN protein was bound to TALON Metal Affinity Resins (Clontech) at 4°C. The gel was washed 3 times with purfication buffer, and eluted using elution buffer [100 mM NaP_i_ (pH 8.0), 200 mM sodium chloride, 150 mM imidazol]. The concentrated fraction was applied to Mono Q 5/50 GL column (GE Healthcare) at 4°C using AKTA Pure HPLC system (GE Healthcare). The column was washed with 20 mM NaP_i_ (pH 8.0) and the sample was eluted using potassium chloride gradient. High concentration fraction was mixed with glycerol (final concentration 15∼20%), aliquoted, frozen by liquid nitrogen and stored at -80°C. The protein concentration was determined by absorbance at 280 nm, and converted to concentrations using molecular extinction coefficient for QUEEN (*ε* = 24,870 M^-1^ cm^-1^, predicted from tryptophan and tyrosine residue numbers).

### DNA constructs for mammalian expression

In order to express QUEEN-37C in mammalian cells, QUEEN-37C was first cloned into pN1/EGFP vector together with Kozak sequence (ACCATGGTG) at the translation initiation site and obtained pN1/QUEEN-37C. Next, to stably transform cells using Tol2 transposon gene transfer system (Kawakami et al, 2007), this QUEEN sequence of pN1/QUEEN-37C was further cloned into a vector containing Tol2 transposon sequences and we obtained pT2MP.EF-QUE37C (#129332). This vector also contains EF1*α* promoter for target protein expression (Addgene #65712) and puromycin resistance gene under PGK promoter for antibiotic selection of the transferred cells. The original vector for this was created and kindly provided by Dr. Akira Takai.

To stably express QUEEN-37C using lentiviral transfection, pN1/QUEEN-37C sequence was cloned into pLBS lentiviral transfer vector together with either EF1*α* or CAG promoter, and we obtained pLBS-EF-QUE37C and QUE37C/pLBS.CAG (Addgene #129338 and #129340). The original vectors pLBS.EF and pLBS.CAG (Addgene #129335 and #129349) were created and kindly provided by Dr. Daisuke Ino.

For expression of ratiometric pHluorin (Miesenbock et al, 1998) in cytosol, pHluorin sequence (a gift from Dr. Hiromi Imamura) was cloned into pN1/EGFP vector, and then into the Tol2 transposon vector and we obtained pT2MP.EF-pHlu (Addgene #129304). For nuclear localization of pHluorin, nuclear localization signal sequence (amino acid sequence PKKKRKV) was fused to the N-terminus of pHluorin by PCR, and then cloned into lentiviral transfer vector to obtain pLBS.EF_NLS-pHlu (Addgene #129324). For expression of FUCCI markers, mScarletI (Addgene #85044, a gift from Dorus Gadella) was fused on the N-terminus of hCdt1 (Addgene #80007, a gift from Vladislav V. Verkhusha). miRFP709-hGem was a gift from Vladislav V. Verkhusha (Addgene #80006). These FUCCI proteins were cloned into pLBS plasmid carrying EF1α promoter, and we obtained pLBS.EF/miRFP670-hGem110Nt and pLBS.EF/mScarlet(i)-hCdt100Nt (Addgene #129333 and #129334). For the expression of cell lineage marker in mES cells, mScarlet (Addgene #85042, a gift from Dorus Gadella) and iRFP713 (a gift from Michiyuki Matsuda) was cloned into pLBS vector and thus we obtained pLBS-mScarlet (Addgene #129337) and pLBS.EF-iRFP (Addgene #129329). For expression of G3BP1-iRFP, human G3BP1 (cloned by Dr. Akira Takai) and iRFP713 (a gift from Dr. Michiyuki Matsuda) was cloned into pN1 vector and we obtained pN1/G3BP1-iRFP (Addgene #129339).

### Lentivirus preparation

To produce lentivirus, HEK-293T cells were transformed with pLBS transfer vector containing DNA sequence of protein of interest, together with envelope/packaging vectors pMD2.G and psPAX2. pMD2.G and psPAX2 was a gift from Didier Trono (AddGene #12259 and #12260). 24 h later, the culture medium was exchanged to Nb4 medium. After additional 24 h, the culture supernatant was collected and concentrated using the Amicon Ultra centrifugal filter unit with 100 kDa membrane (Merck), aliquoted and stored in -80°C.

### In vitro fluorescence measurements

For in vitro measurements, 10-30 μl of ∼5 μM QUEEN was diluted in 2000 μl measurement buffer [50 mM HEPES-KOH (pH 7.3), 50 mM potassium chloride, 1 mM magnesium sulfate, 0.05 % Triton X-100] in a polystyrene cuvette. Fluorescence excitation spectrum was measured using a spectrofluorometer (JASCO FP-8200 for QUEEN-37C and JASCO FP-6200 for QUEEN-20C) at 513 nm emission. Typically, the spectra showed two peaks at 400 nm and 494 nm excitation. MgATP solution (equimolar mixture of ATP and magnesium sulfate) was added to cuvette and measurement was repeated. Temperature was checked before each measurement and difference from the target temperature was always no more than ± 0.3°C. The ratio of intensity at 400 nm and 494 nm excitation (*R*) was plotted against ATP concentration (*c*), and the data points were fitted to Hill equation,

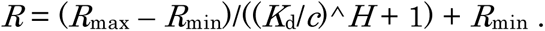

*H* is the hill coefficient, and *K*_d_ is the dissociation constant of QUEEN to ATP. *R*_max_ and *R*_min_ are the maximum and minimum ratio values.

For measurement at pH 7.6 and 7.9, measurement buffer component was the same except for pH. For measurement at pH 7.0 and 6.7, HEPES was replaced with PIPES and the pH was adjusted to these values.

For kinetic measurements, 10-15 μl of ∼5 μM QUEEN-37C sample was diluted in 1500 μl measurement buffer in cuvette and kept at 37°C. MgATP solution was mixed with measurement buffer to make 500 μl of 4× ATP stock. This ATP stock solution was rapidly added to the cuvette through a thin silicone tube and mixed. ATP solution was pre-heated to ∼50°C prior to mixing, so that the sample temperature was kept at 37°C after liquid transfer. Fluorescence timecourse at 488 nm excitation and 513 nm emission was measured to observe the relaxation of the system to new equilibrium. To omit the effect of intensity change by sample dilution, only the last 15% (based on fluorescence intensity) of the fluorescence change was used for fitting to single exponential curve. The inverse of time constant was considered as the rate of relaxation to the new equilibrium. k_on_ and k_off_ was obtained as the slope and y-intercept of the linear fitting of the plot. Fitting was done using Kaleidagraph (Synergy Software).

### Cell culture

All the cells were cultured at 37°C under 5% CO_2_ atmosphere. MDCK and HEK293T cells were cultured in DMEM 10% FBS [High glucose DMEM (Wako #044-29765), supplemented with 10% Fetal bovine serum (Gibco #10270-106), 100 U/ml penicillin and streptomycin (Gibco #15140163)].

For the primary hippocampal neuron and astrocyte culture, glass-bottom dishes were first coated with 0.04% polyethyleneimine for 2 h and 1 mg/ml poly-L-lysine for 2h. MEM 10%HS [MEM (Gibco #11090-081) supplemented with 10% horse serum (Gibco 16050-122), 33 mM Glucose, 1 mM pyruvate, 2 mM GlutaMax, 100 U/ml penicillin and 100 μg/ml streptomycin] was added to the dish and equilibrated to 5% CO_2_ overnight. Hippocampus was dissected from mouse embryo, and enzymatically dissociated by papaine (Worthington) for 20 min and DNaseI (Roche) for 5 min. Then the cells were mixed with the medium in the coated glass-bottom dishes. After the cells were attached to the glass, culture medium was exchanged to Nb4+MSG [NbActiv4 medium (BrainBits) supplemented with 25 μM monosodium glutamate (MP Biomedicals)]. After 4 days, half of the medium was exchanged with NbActiv4 medium.

For the mouse embryonic fibroblast culture, lung was dissected from mouse embryo, minced and plated on uncoated polystyrene dishes. DMEM 10% FBS was used as the culture medium. For mES cells (EB5), cells were cultured in GMEM (Gibco #11710-035) supplemented with 10% knockout serum replacement (Gibco #10828-028, 1%FBS, 1 mM sodium pyruvate (Gibco #11360-070), 1x non-essential amino acids (NEAA, Gibco #11140-050), 0.1 mM 2-mercaptoethanol (Sigma-Aldrich) and 2,000U/mL leukemia inhibitory factor (Nacalai Tesque NU0013-1). The cells were plated on gelatin-coated polystyrene or glass-bottom dishes.

For the culture of other cells, culture was done according to the conditions recommended by the cell provider was used. The medium and culture conditions are summarized in Supplementary Table S3. DMEM, Low glucose DMEM (DMEM(LG), #041-29775), EMEM (#051-07615), HamF12 (#087-08335), RPMI1640 (#189-02025), DMEM/Ham’s F-12 (#042-30555) and Hygromycin B were purchased from Wako Pure Chemicals. Endothelial Cell Growth Medium 2 (C-22111) was purchased from PromoCell. McCoy’s 5a (Gibco #16600082) was purchased from Thermo Fisher.

### Transformation of mammalian cells

MDCK cells stably expressing QUEEN-37C were obtained by transfecting MDCK II cells with pT2MP.EF-QUE37C vector together with pCS-TP transposase vector (Kawakami et al, 2007) using TransFectin Lipid Agent (Bio-RAD). The transfected cells were cultured under 1 μg/ml puromycin condition, and a fluorescent colony was isolated to obtain a cell line that stably express QUEEN-37C. MDCK cells stably expressing cytosolic pHluorin was transformed into MDCK cells in a similar way using pT2MP.EF-pHlu vector.

All the cells in Table 1 was transfected with lentivirus produced from either pLBS-EF-QUE37C or QUE37C/pLBS.CAG. pLBS-EF-QUE37C lentivirus was used for A549, C2C12, CaCo-2, HCT116, L929 and NIH3T3 cells. QUE37C/pLBS.CAG lentivirus was used for the rest of the cells in Table 1. We did not selectively isolate the fluorescent cells for these experiments. The cells were cultured at least 3 days after transfection before imaging.

For the simultaneous measurement of ATP and pH, MDCK was transformed with pT2MP.EF-QUE37C first. The cells with relatively weak fluorescence intensity were isolated and propagated. This line is hereafter referred to as Q37_Low_ cells. Then Q37_Low_ cells were transfected with lentivirus from pLBS.EF_NLS-pHlu and cells with strong nuclear fluorescence were isolated again. This line is hereafter referred to as Q37Low/nuc-pHlu cells. For the cells expressing nuclear pHluorin alone, wild type MDCK cells were transfected with lentivirus produced from pLBS.EF_NLS-pHlu.

For the simultaneous measurement of ATP and cell cycle, MDCK cells expressing QUEEN-37C were transformed with lentivirus produced from pLBS.EF/miRFP670-hGem110Nt followed by lentivirus produced from pLBS.EF/mScarlet(i)-hCdt100Nt. A single colony with good expression levels of both proteins was selected and propagated.

For measurement of correlation within the colony of mES cells, EB5 mES cells were transformed by lentivirus produced from pLBS-EF-QUE37C. Then, these cells were transformed by lentivirus produced from either pLBS-mScarlet or pLBS.EF-iRFP. The cells with good levels of fluorescence were selected using flow cytometry (FACS Aria III, BD Biosciences).

For measurement of SG formation, HEK293T cells were transfected with lentivirus produced from QUE37C/pLBS.CAG vector. Then, cells were transiently transfected with pN1/G3BP1-iRFP plasmid using TransFectin Lipid Agent.

We deposited the stable cell lines created in our study to RIKEN Cell Bank to facilitate the usage of other researchers in the field (Supplementary Table S4).

### ATP quantification by luciferase

MDCK cells expressing QUEEN-37C were dissociated from the plate by incubating in TrypLE Express at 37°C for 15 minutes. After mixing with twice the volume of DMEM 10%FBS, the cells were centrifuged and suspended in DMEM 10%FBS conditioned by MDCK cells. The cells were imaged first using A1R confocal microscope (see the next section for imaging conditions). Next, 160 μl luciferase sampling buffer [100 mM Tris-HCl (pH 7.75) and 4 mM EDTA] and 20 μl spike ATP solution (equimolar mixture of 0 – 1000 nM ATP and magnesium sulfate) were mixed and pre-heated at 95°C. 20 μl of cell suspension was transferred to this pre-heated mixture, kept on 95°C for 10 min to denature proteins, placed on ice and stored at -80°C. These samples were assayed by ATP Bioluminescence Assay Kit CLS II (Roche) afterwards. Luminescence was quantified using microplate reader MTP-880 (Corona Electric).

The measured values were plotted against spike ATP concentration. As the spike ATP was increased, the measured ATP values increased too, but the slope of linear regression was always significantly smaller than 1 (typically ∼0.7). We surmised that this is probably because ∼30% of ATP was consumed by enzymes inside cells during heat inactivation. Intracellular enzymes might have remained active for a short time after cell rupture by heat, before they were completely inactivated. Assuming that a constant fraction of ATP is consumed, ATP concentration of the heat-inactivated sample (*C*) is related to ATP concentration from the cytosol if no ATP consumption occured (*X*) as in,

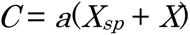

where *a* is the slope, and *X_sp_* is the final concentration of added spike ATP. *X* can be obtained from linear regression of the *C*-*X_sp_* plot as the absolute value of *X_sp_* axis intercept of the regression line. *X* is related to the ATP concentration in cells as in,

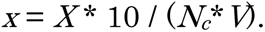

where *N_c_* is the number of cell per cell suspension volume and *V* is the volume per single cell. 10 is to correct for the dilution occurring in the sampling procedure.

The number of cells per 1 ml suspension was counted using Neubauer hemocytometer chambers (ISOLAB). Four 1 mm x 1 mm square region (0.1 mm in height, 0.1 µl in volume) of the chamber was counted, and average was calculated. The average cell volume of each sample was determined from the z stack image of the 488 nm excitation channel (see below). Estimation error of *X* was calculated as,

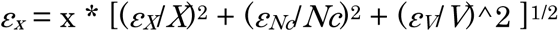

where *ε_X_*, *ε_N_* and *ε_V_* are the standard error of mean values of *X*, *N_c_* and *V*.

### Fluorescence imaging

Cells were plated on glass-bottom dish (MatTek P35G-0-14-C/H) coated with CellMatrix Type I-C (Nitta Gelatin), unless otherwise stated. For the imaging of MDCK cells expressing QUEEN-37C, cells were cultured for 1-4 days after plating until they reach the desired CD. For imaging of MDCK cells expressing FUCCI cell cycle reporters, cells were cultured for 1-2 days after plating. For all the cells in Table 1, cells were cultured for 2-4 days after plating. For imaging of mES cell colonies, the glass bottom dish was coated with Matrigel (Corning), and cells were cultured 24 h after plating. Images were acquired using Nikon A1R confocal microscope system. The microscope unit Ti-2 was equipped with 40×/1.25 WI λS objective lens. For imaging of QUEEN-37C, two images were taken, namely at 405 nm excitation (405ex) and 488 nm excitation (488ex). Light source for 405ex was 1170506/AC (COHERENT, light source 405). Two different light sources were used for 488ex throughout this study, namely IMA101040ALS (Melles Glliot, light source 488-1) and LGK7873ML (LASOS, light source 488-2). For the experiments in Fig. 2, 405ex and 488ex channels were acquired alternately for each line (“line-by-line” mode). Light source 488-1 was used for these measurements. For the experiments in Table 1, Figure 3-6 and Supplementary Fig S2-3, 405ex and 488ex channels were acquired one image at a time (“image-by-image” mode). Light source 488-2 was used for these ones. Emission light was collected through 525/50 emission filters. In all cases, acquisition image size was 1024 × 1024, which corresponds to 318 × 318 µm^2^ area. Excitation laser intensity was estimated to be ∼1 µW, and image acquisition rate was 0.5 frame per second (pixel dwell time was 1.1 µs / pixel). 4× averaging was applied. During all measurements, temperature was controlled to 37 ± 0.5 °C and CO_2_ was controlled to 5% using a stage top incubator (Tokai Hit), unless otherwise stated.

For size estimation of detached round cells in luciferase assay, z-stack was acquired using the 488ex channel with 1 µm steps. For size measurement using DiD membrane staining, MDCK cells expressing QUEEN-37C were stained in medium containing 5 µg/ml DiD solution for approximately 10 minutes. Light sources used were light source 488-1 for 488 nm and 1170508/AC (COHERENT) for 640 nm (light source 640). For 2-DG and oliA treatment of detached cells, cells were removed from the plate and collected by centrifugation, and resuspended in DMEM medium containing the drugs of desired concentrations. The suspension was put in glass-bottom dish and was incubated for ∼10 minutes in 37°C 5% CO_2_ until many cells settle to the bottom of the dish before imaging. We did not equilibrate the DMEM in CO_2_ in advance. In such cases, the intracellular pH showed high values (>8.0) right after medium exchange, but under 5% CO_2_ it soon reached within the range of 7.0-8.0 in 10 minutes (Supplementary Fig. S2C), so the effect of the non-CO_2_-equilibrated medium to the measured ATP value in Fig. 2 is likely to be small. We also tested if normal DMEM and no riboflavin DMEM show different results. Cells grown in no riboflavin DMEM were detached and measured similarly. It turned out that no riboflavin DMEM culture showed similar results to normal DMEM culture, so these two groups were combined and plotted as “No Drug”.

For oligomycin A treatment of neurons and astrocytes, oliA was directly added to the dish to final concentration of 10 µg/ml. For immunolabeling of neurons and astrocytes, cells were washed with PBS, fixed with 4% paraformaldehyde and permeabilized with 0.3% Triton-X. Cells were incubated with 3% BSA, and then with 3% BSA, 1% mouse anti-MAP2 (MS X MAP-2, Millipore) and 0.1% rabbit anti-GFAP (GFAP-Rb-Af800, Frontier Bioinstitute). Then the cells were labeled with 3% BSA, 10 µg/ml goat Alexa568-anti mouse and 10 µg/ml goat Alexa647-anti rabbit (Thermo Fisher). Cells were imaged using light source 488-2 for 488 nm and SAPPHIRE 561-20 CW CDRH (COHERENT, light source 561) for 561 nm excitation.

For oxa and oliA treatment of attached MDCK cells, cells were plated on glass-bottom dish coated with collagen and was cultured until it reached the desired cell density. Then the culture dish was placed on the microscope and silicone tubes and a temperature probe were attached. The tubes are used for exchanging medium in the dishes. As a mock treatment, old medium was exchanged to “mock” medium (DMEM 10%FBS medium with 20 mM mannitol and 0.1 % DMSO). Subsequently, the medium was exchanged to either “oxamate” medium (medium with 20 mM oxamate and 0.1% DMSO), “oliA” medium (medium with 20 mM mannitol and 10 µg/ml oliA) or “oxamate&oliA” medium (medium with 20 mM oxamate and 10 µg/ml oliA), depending on the experiment. The new medium was equilibrated in advance to 5% CO_2_ atmosphere to adjust pH. Interval between frames was 70 seconds (Fig.4A-D) or 20 minutes (Fig. 4E-F).

For pHluorin calibration, cyt-pHlu or Q37_Low_/nuc-pHlu cells were incubated in nigericin medium (DMEM 10% FBS containing 10 µM nigericin amd 10 µM valinomycin) for 30 minutes. Then, the CO_2_ control was stopped, and the extracellular medium was exchanged to the buffer of various pH [20 mM HEPES (pH ≥ 7.3) or PIPES (pH ≤ 7.0), 120 mM potassium chloride, 1 mM magnesium sulfate, 2 mM calcium chloride, 10 mM Glucose] sequentially. Images were acquired every 70 seconds.

For FUCCI measurement in MDCK cells, and for mScarlet and iRFP imaging in EB5 mES cells, light source 561 and light source 640 was used for excitation.

For measurement of SG formation, the imaging was performed 2 days after pN1/G3BP1-iRFP transfection. iRFP was excited using light source 640. Arsenite (50 mM sodium arsenite solution, Merck-Millipore) was added to the dish during imaging at a final concentration of 100 µM. OliA and oxamate treatment was done in the same way as the MDCK cells.

### Image processing for cell size analysis

For cell size estimation in luciferase assay, semiautomatic segmentation of round-shaped cells was used to estimate the average cell size. 1024 × 1024 image was used without resizing. Using fluorescent beads of uniform diameter (Thermo Fisher F21011), we confirmed that length calibration in our microscope system was accurate. In principle, thresholding the spherical cell image by Li’s algorithm (included in ImageJ) provided a quite good estimation of the cell border. We experimentally determined the precise membrane position using DiD membrane staining dye (Thermo Fisher, D7757). We found that after thresholding the 488ex channel image by Li’s algorithm and then eroding the binary mask by 3 px, the mask border matches well with the true membrane position. However, because QUEEN-37C-expressing cells have nuclei that appear as low intensity blobs in the fluorescence image, simple usage of Li’s algorithm will leave holes inside cell masks. This greatly hampers the automatic separation of touching cells by watershed. To remove such holes, “triangle mask” image (thresholded with triangle algorithm) was subtracted from “Li mask” (thresholded with Li’s algorithm) image, and then morphological opening (2 px) was applied. This “nuclei blob” image was subtracted from the “Li mask” image and morphological opening (2 px) was applied. This resulted in a “blob-filled” image with most of blobs filled. The “blob-filled” image was inverted, and watershed was applied to separate the touching cells to make “separated cells” image. Finally, morphological erosion (3 px) was applied to make “cell size” image. The area of each particle was measured and exported as a text file for size calculation. This procedure was repeated for all the slices in the z-stack of 1 µm step, and the largest area value for each cell (*S_i,max_*) was used to estimate the volume (*V_i_)* of cell *i*.

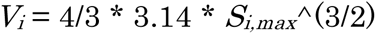

Then, population average and standard deviation for *V_i_* was calculated.

### Shading correction

The QUEEN-37C ratio images are just for visualization and shading correction was not applied during creation of these images. For the quantitative analysis in Table 1, Fig. 3-6 and Supplementary Fig. S2, shading correction was applied before analysis. Assuming that x is the position in the image, the raw image *I*(x) is related to the true expected signal *R*(x) by

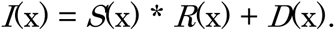

*S*(x) is the shading of the illumination field, and *D*(x) is the baseline of the camera. In our confocal imaging conditions, background signal from medium was very low and could be ignored. Therefore, *D*(x) can be substituted with a constant value, *D*_b_, which is the mean intensity of the background region. To obtain *S*(x), a small piece of cover slip was fixed on top of the 35 mm glass-bottom dish using a 10 µm-thick adhesive tape. The thin gap between the glass was filled with 1 µg/ml fluorescein solution. This was imaged under A1R confocal microscope using almost the same imaging conditions as the cells, except that the stronger laser intensity for 405 nm excitation was used to compensate for the weak excitation efficiency of fluorescein at 405 nm. 15-20 images were taken at different XY positions and Z-projected (median) using ImageJ/Fiji (Schindelin et al, 2012) to remove position-specific artefacts. This Z-projection image had weaker intensity at the edges. Then, the intensity at the center 60 × 60 pixel ratio of the 512 × 512 image was normalized to 1, and median (20 px) and mean (20 px) filters were applied for smoothing. This normalized image was considered as *S*(x). Shading-corrected image of the cells were thus obtained by,

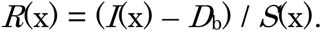

In practice, zero or negative values are undesired for further analysis. Thus, we added *D*_b_ back to *R*(x) to create corrected image used for analysis, *R*_a_(x).

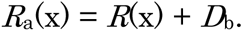

### Image processing for ratio calculation

All the acquired fluorescence images were first resized from 1024 × 1024 to 512 × 512 in NIS Elements software (Nikon) before further image processing. Ratio color images were created using NIS Elements.

To analyze the ratiometric sensor images, briefly, 405ex and 488ex images were shading-corrected, regions of interest (ROI) were created inside single cells, ratio value was calculated for each pixel inside ROI and aggregated for all the pixels in the ROI. Shading correction and ROI creation were done using ImageJ/Fiji software and pixel values were exported as text files for further analysis in Python 3.7.

In some experiments (Fig. 2B-C, 4, 6C-E, Supplementary Fig. S2D-H, S3 and S4), ROIs were created semi-automatically using ImageJ Macro (see below for details). In other imaging measurements (Fig. 2A, 3, 6A, Supplementary Fig. S1 and S2A-C), ROIs were created manually.

For imaging of QUEEN-37C MDCK cells in Fig. 4, Supplementary Fig. S3 and S4, semi-automatic segmentation was used to find the nuclei, and then the pixels surrounding the nuclei (3 px width) were used as ROIs for cytosol regions. Using our self-made macro, nuclei were extracted from the images with good accuracy, with true positive rate of >80% and false discovery rate of <10%. For timelapse measurements, single-cell positions were tracked using TrackMate plugin (Tinevez et al, 2017) in ImageJ/Fiji.

For imaging of QUEEN-37C mES cells in Fig. 6A, ROIs were created manually in the cytosol region. Colonies including more than 3 cells were used for analysis. For imaging of co-culture of mScarlet or iRFP lines of QUEEN-37C mES cells in Fig. 6C-E, nuclei were semi-automatically detected and pixels surrounding nuclei were used as ROI for cytosol, as in MDCK cells. Only the colonies including more than 3 cells of both cell lines were used for analysis.

For imaging of QUEEN-37C MDCK cells in Fig. 2B-C, rounded cells were detected semi-automatically using ImageJ/Fiji macro. Then, to avoid the nuclei with relatively weak signal intensity, only the edges (3 px width) of each rounded cells were used as ROIs for ratio calculation.

For the measurement of nuc-pHlu and Q37_Low_/nuc-pHlu cells in Supplementary Fig. S2D-H, nuclei were semi-automatically detected using the stronger nuc-pHlu fluorescence. When necessary, pixels surrounding the nuclei were used as cytosol ROI.

Cell number per image field was counted semiautomatically using ImageJ. The number of cell nuclei in the image was used as the cell number per image. Imaging area per image was 0.100 mm^2^ (317 µm x 317 µm). Nuclei on the edge of the image were excluded.

### Ratio, ATP and pH calculation

The exported text files of pixel intensities were analyzed using self-made Python 3.7 program. To calculate the ratio of QUEEN-37C or ratiometric pHluorin, background intensity was subtracted, and the 405ex/488ex ratio for each pixel was calculated. The median value of all the pixels in one ROI was used as the representative value of the cell. Coefficient of variance (CV, SD divided by mean) of ratio was also calculated for each cell ROI, and the cells with ratio CV over a certain value (>0.4 for MDCK cells, >1 for mES cells) were considered to be too noisy and were omitted from data.

For the measurements of ATP using QUEEN-37C, the ratio *R* was further converted to ATP concentration *c* using the inverse of Hill’s equation,

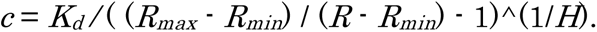

The values obtained in in vitro measurements using spectrofluorometer were used for *K_d_* and *H*. In our measurements, *K_d_* was 2.92 and *H* was 2.24. *R_max_* and *R_min_* values were determined using SLO permeabilization calibration method for each microscopy conditions (see the next section). If *R* > *R_max_*, or if *c* was calculated to be >15 mM, QUEEN-37C was considered to be saturated, and 15 mM was used as the value for *c*. If *R* < *R_min_*, 0 mM was used as the value for *c*. Median ratio in the ROI was converted to ATP concentration, and this was used as the representative ATP concentration for the ROI.

For identification of cell cycle using FUCCI, The ratio of 561 nm excitation divided by the sum of 561 nm and 640 nm excitation was calculated for each cell, and if the value was over 0.6, the cell was considered to be in the G1 phase.

For QUEEN-37C ratio calculation of cell colonies with mScarlet or iRFP cell lines, intensity of 561 nm and 640 nm excitation was used to separate the two cell lines. If both 561 nm and 640 nm excitation signals were detected in the same pixel, the pixel was not used so as to omit the overlapped region of two different cell types.

For pH and ATP simultaneous measurements in Q37_Low_/nuc-pHlu cells, nucleus fluorescence (mainly from nuc-pHlu) and cytosol fluorescence (mainly from QUEEN-37C) was separately measured for each cell. In brief, the strong nucleus fluorescence intensity of nuc-pHlu was determined first, and then the weak fluorescence of cytosol QUEEN-37C was corrected for nuc-pHlu leakage and its intensity was determined. Q37_Low_/nuc-pHlu cells were thresholded by nuc-pHlu intensity, so that they had more than 10 times stronger fluorescence in the nuclei than the nucleus region of Q37_Low_ cells (Supplementary Fig. S2D). Therefore, the fluorescence from QUEEN-37C in nuclei can be ignored in practice for our Q37_Low_/nuc-pHlu cells.

To obtain the pHluorin pH calibration curve from nigericin/valinomycin experiment, mean ratio values of Q37_Low_/nuc-pHlu cells were plotted against pH and fitted to a first-degree polynominal to obtain a linear calibration curve (Supplementary Fig. S2A). This calibration curve was used to convert ratio to pH values for each pixel, and the median value in each ROI was plotted.

To correct for the nuc-pHlu leakage in the cytosol of Q37_Low_/nuc-pHlu cells, the cytosolic intensity of nuc-pHlu cells (i.e. no QUEEN-37C expression) was measured. We found that the intensity of nucleus and cytosol in nuc-pHlu cells were linearly correlated. This correlation was used to estimate the nuc-pHlu leakage in the cytosol of nuc-pHlu/QUEEN-37C cells using nucleus nuc-pHlu fluorescence (Supplementary Fig. S2E). The leakage-corrected ratio for QUEEN-37C was calculated for each pixel and median of the cytosol pixels was determined. The level of estimation error was calculated by bootstrapping. Median ratio, and its 95% confidence interval values, was converted to ATP values as above.

### Estimation of cell area by voronoi segmentation

To estimate the single-cell area in Supplementary Fig. S3, nuclei was semi-automatically detected first using ImageJ/Fiji macro. Then, the binarized image of nuclei was used as the seed for Voronoi function of ImageJ/Fiji. The area of Voronoi-segmented regions were used as the estimates of single-cell area in the image. Segmented regions touching the edge of the image was omitted from analysis.

### Creating plots and statistical analysis

The time traces of many single cells in Fig. 4 and Supplementary Figure S2G-H was created using Python 3.7. Time traces with missing value gap larger than 5 frames were excluded from the plot. 8-20 representative time traces are highlighted in different colors for visualization, and all the other time traces are shown in gray.

Scatter plots in Supplementary Fig. S3, and strip plots in Fig. 6 and S3 were created using Python 3.7.

One-way ANOVA (Fig. 6A, Supplementary Fig. S3E-F), Python 3.7’s HSD post hoc test (Supplementary Fig. S3E-F), two-way ANOVA (Table 2) and Student’s t-Test (Fig. 6E) was performed using RStudio software.

All the other line plots, scatter plots and box-whisker plots of ATP and pH values in this report were created using Kaleidagraph.

### Calibration of QUEEN signal by cell permeabilization

The absolute *R_max_* and *R_min_* values of 405ex/488ex ratio are dependent on measurement setups. Accordingly, to convert the change of 405ex/488ex ratio obtained by microscope imaging to actual ATP values, we needed to calibrate for the difference between spectrofluorometer and microscope conditions. For this purpose, we performed cell permeabilization experiments using streptolysin O (SLO) (S5265, Sigma-Aldrich). MDCK cells expressing QUEEN-37C was plated on a 35-mm glass bottom dish and cultured to ∼80% confluency. Calibration buffer [50 mM HEPES-KOH (pH 7.3), 50 mM potassium chloride, 1 mM magnesium sulfate] was prepared in advance. The cells were washed 1-3 times with calibration buffer and 160 µl of calibration buffer was added to the center glass area of the dish. ∼20 µl MgATP stock solution (200-300 mM equimolar mixture of ATP and magnesium sulfate) was added at a final concentration of 20 mM for saturating ATP conditions. MgATP was not added for 0 mM ATP conditions. 10 µl of 25,000 U/ml SLO stock was then added to the dish and mixed by gentle pipetting. After the temperature has returned to 37 °C, 405ex and 488ex fluorescence images were acquired. Two different acquisition modes were used (see “Fluorescence imaging” section). For the experiments in Fig 2, two channels were acquired alternately for each line (“line-by-line” mode). Interval between frames was 60 sec. For the other experiments using QUEEN or pHluorin under the microscope, two channels were acquired one image at a time (“image-by-image” mode). The time lag between two channels was 10 sec, from midpoint of 405ex to midpoint of 488ex acquisition. Interval between frames was 66 sec. After a few minutes, a fraction of cells showed a rapid decrease in fluorescence intensity, due to QUEEN-37C leakage from formed SLO pores.

We analyzed the intensity timecourse using ImageJ/Fiji and self-made Python 3.7 program. The frame that many cells started to show decrease in their intensity was used for analysis. The ratio of intensity change (intensity in the next frame divided by intensity in the current frame) was calculated for each ROI and channel. These ratios are hereafter referred to as 405Nx and 488Nx for 405ex and 488ex channels, respectively. For line-by-line mode acquisition data, cells that showed 405Nx and 488Nx values of 0-0.9 was used for analysis. For image-by-image mode acquisition data, cells that showed 405Nx and 488Nx values of 0.7-0.9 was used for analysis. Calculation of ratio values was done as stated in the “Ratio, ATP and pH calculation” section. The ratio for each pixel was calculated and median was used as the representative value for that ROI. If the coefficient of variance (CV) of ratio in the ROI was larger then 0.4, that ROI was considered to be too noisy and was omitted. The mean of ROIs in that condition was calculated. For line-by-line acquisition data, these values (2.11 and 0.62, Supplementary Dataset S1) were directly used as either the *R_max_* (20 mM ATP condition) or *R_min_* (0 mM ATP condition) values of the QUEEN-37C calibration curve. For image-by-image mode data, we took into consider the time lag between two acquisition channels. There was a 10 s time lag between the 405ex and 488ex channels, the timelapse interval was 66 s and we analyzed the data points with 405Nx and 488Nx values in the range of 0.7-0.9. Accordingly, to compensate for the intensity change during the time lag, we multiplied 0.967 [= 0.8 ^ (10/66)] to the ratio mean values. These corrected values (1.51 × 0.967 and 0.43 × 0.967, Supplementary Dataset S2) were used as *R_max_* and *R_min_* for the calculation of data taken in image-by-image mode .

### pH correction of ATP concentration

To correct the ATP concentration for pH in nuc-pHlu/QUEEN-37C cells, the relationship of pH and ATP in Fig. 1C was used. Since the ratio for microscopy experiments and spectrofluorometer experiments are not same, we first converted the ratio value in microscopy condition to that in spectrofluorometer condition. First, assuming that the pH was 7.3, an uncorrected ATP concentration value *c_7.3_* was obtained using calibration curve for microscopy conditions. Next, we calculated the ratio value for spectrofluorometer condition that gives ATP concentration of *c_7.3_* at pH 7.3. Hereafter we call this ratio value *r_F_*. In addition, we will refer to the spectrofluorometer condition ratio value when pH is *p* and ATP concentration is *x* as *R*(*p*, *x*), and will refer to the ATP concentration at pH *p* and spectrofluorometer condition ratio value *r* as *c*(p, r).

From the Hill equation fitting in Fig. 1C, *R*(*p_i_*, 0) (*p_i_* : 6.7, 7.0, 7.3, 7.6, 7.9) is known. By fitting *R*(*p*_i_, 0) vs *p* plot to second degree polynominal, we calculated *p* = *p_B_* that satisfies *R*(*p_B_*,0) = *r_F_*. If *p_B_* < 6.7, or in other words, if *r_F_* is larger than *R*(6.7, 0), we can use all five points in *c*(*p_i_*, *r_F_*) (*p_i_* : 6.7, 7.0, 7.3, 7.6, 7.9) to get the ATP-pH relationship with *r* fixed at *r* = *r_F_*. The curve shape of *c*(*p*, *r_F_*) vs *p* plot can be obtained by fitting the points to second degree polynominal. If *p_B_* > 6.7, or in other words, if *R*(6.7, 0) < *r_F_* < *R*(7.9, 0), points in *c*(*p_i_*, *r_F_*) (*p_i_* : 6.7, 7.0, 7.3, 7.6, 7.9) that does not satisfy *p_i_* >*p_B_* was omitted before fitting. In addition, we know that *c*(*p_B_*, *r_F_*) = 0 so we can use it for fitting. The curve of *c*(*p*, *r_F_*) vs *p* plot was obtained by fitting the available data points to Hill equation. Once we obtained the relation between *c*(*p*, *r*_F_) and *p*, we can calculate the corrected ATP concentration by *c*(*p_MS_*, *r_F_*), where *p_MS_* is the pH value measured by nuc-pHlu imaging.

### Calculation of spatial autocorrelation

To calculate autocorrelation of ATP concentration without any shuffling, the ratio of 405ex and 488ex channels were calculated for each pixel. Next, to minimize the effect of noise, the 512 × 512 original image was resized to 64 × 64 coarse-grained image. This was done by gridding the original image to 64 × 64 regions, and the median in each region was used as the value of the corresponding pixel in the coarse-grained image. Then, the ratio values in each pixel of the coarse-grained image were converted to ATP values. Finally, the autocorrelation of this ATP value image was calculated. The autocorrelation values were radially averaged. We used a macro written by M. Schmid for the radial autocorrelation calculation step (http://imagejdocu.tudor.lu/doku.php?id=macro:radially_averaged_autocorrelation).

For the simulated image of scrambled pixel values within each cells, we first needed to determine the position of each cells. To do this, the positions of nuclei were detected, and this was used as seeds for watershed segmentation to create a segmented image. Each compartmentalized area corresponds to a region occupied by a single cell, so we hereafter refer to these regions as “pseudo-cells.” Borderline pixels were merged into one of the neighboring pseudo-cells so that every pixel belongs to one of the pseudo-cells. Then, one pseudo-cell region was selected, and all the pixels in this region were scrambled. Subsequently, image was resized to 64 × 64, ratio values were converted to ATP values, and the radial autocorrelation was calculated in the same way as above. Although the simple segmentation procedure we used was not perfect and some inconsistency from the real data was present, this did not effect the final autocorrelation values of the coarse-grained images (Fig. 5I, J).

For the simulated image of scrambled cell position, one pseudo-cell region was chosen, and the ratio values from a second pseudo-cell, which was chosen irrelevantly from the first one, was used to fill inside the first pseudo-cell. The rest of the procedure is the same as scrambling within each cell.

---

Supplementary Fig. S1

Simultaneous measurement of ATP concentration and G3BP1 foci formation. HEK293T cells expressing G3BP1-iRFP and QUEEN-37C was treated by arsenite (A, C) or oliA and oxamate (oliA&oxa) (B, D). (A, B) Representative images of QUEEN-37C ratio (top) and G3BP1-iRFP fluorescence (bottom) during arsenite (A) or oliA&oxa (B) treatment. Approximate time after measurement start is shown in the corner in minutes. Scale bar = 10 µm. (C, D) Time courses of ATP concentration (top) or CV (coefficient of variance) of G3BP1-iRFP fluorescence (bottom) during arsenite (C) or oliA&oxa (D) treatment. Each trace corresponds to single cells. For clarity, 8-10 traces are highlighed in color solid lines, and the top and bottom panels share the color codes. The trace with dashed line and open circles corresponds to the representative cell in (A) or (B). Other traces are shown in gray.

Supplementary Figure S2, related to Figure 4

Measurement of pH during inhibitor treatment. (A) pHluorin calibration using cytosolic QUEEN-37C & nucleic pHluorin (Q37_Low_/nuc-pHlu) cells. Error bar = SD. (B) Box-whisker plots of nucleic pH before and after ATP synthesis inhibitor treatment in nucleic pHluorin (nuc-pHlu) cells. (C) Timecourse of pH in cells expressing cytosolic pHluorin (cyt-pHlu) and nuc-pHlu cells during inhibitor treatment. The two kinds of cells were cultured and measured together in the same dish. Cells were treated by mock and oxamate-containing DMEM medium at indicated points. The medium was not equilibrated in advance to 5% CO_2_ in this experiment. Error bar = SD. (D) Fluorescence intensity in the nucleus region of the cells expressing either QUEEN-37C only (Q37_Low_ cells, blue crosses) or QUEEN-37C and nuclear pHluorin (Q37_Low_/nuc-pHlu cells, red and pink circles). For analysis of Q37_Low_/nuc-pHlu cells, cells with nuclear pHluorin intensities above a certain threshold (red circles) were used so that QUEEN-37C signal in the nucleus can be ignored. (E, top) Fluorescence intensity of the nuclear and cytoplasmic regions of cells expressing nuclear pHluorin. (E, bottom) Total fluorescence intensity of the cytoplasmic region of Q37_Low_/nuc-pHlu cells cells (red circles), and the estimated intensity of nuc-pHlu fluorescence included in it (blue crosses). (F-H) Simultaneous measurement of ATP (cytosolic QUEEN-37C) and pH (nucleic pHluorin) of MDCK cells. (F) Low CD cells were treated by oxamate, and representative trace of a single cell with relatively large pH change is shown. Uncorrected ATP trace (blue) and pH-corrected ATP trace (red) are shown. Broken and dash-dot lines indicate the lower and upper 95% confidence interval of the ATP value estimation. (G, H) Timecourses of pH (top) and pH-corrected ATP (bottom) for a number of cells. Each single trace represents one cell, and 10 of them are highlighted in colors for clarity. (G) Low CD cells were treated by oxamate,. (H) high CD cells were treated by oliA.

Supplementary Figure S3, related to Figure 4

Correlation between apparent cell area and single-cell ATP concentration after metabolism inhibitor treatment in MDCK cells at 3500-4500 cells/mm^2^. (A, B) Example ratio images after voronoi segmentation of cells. Positions of nuceli were used as seeds of voronoi segmentation. The area of segmented regions was used as the estimate values of cell area. (C, D) Scatter plots and its marginal distribution of single-cell area and ATP concentration. (E, F) Single-cell ATP concentration grouped by rank of cell area. The alphabets are compact letter display of the significantly different (p > 0.05) groups found by Tukey’s HSD test. (A, C, E) 10 minutes after oxamate treatment. (B, D, F) 10 minutes after oliA treatment.

Supplementary Figure S4, related to Fig. 4

Response to oliA and oxamate treatment at various cell density. (A) Cells were treated with oliA first, and then with both drugs. (B) Cells were treated with oxamate first, and the with both drugs. to ATP.

Supplementary Table S1

QUEEN candidate variants created in this study and their affinity

Suppelemntray Table S2

List of various constructs created in this study and their Addgene deposit numbers. Fig. 3.

Supplementary Table S3

List of culture conditions for the various cells shown in Table 1 and

Suppelemntray Table S4

List of stable mammalian cell lines created in this study. These cells are available from RIKEN BioResource Research Center upon request.

Supplementary Movie S1

Movie for timecourse measurement of QUEEN-37C ratio values in Fig. 4A-D were concatenated in one movie. Time after starting the measurement is shown on right top corner in minutes.

Supplementary Movie S2

Movie for selected image field of QUEEN-37C measurement during oliA test in Fig. 4E. Time after starting the measurement is shown on right top corner in minutes. One field was selected for each cell density and drug conditions.

Supplementary Movie S3

Movie for selected image field of QUEEN-37C measurement during oxamate test in Fig. 4F.

Supplementary Dataset S1

Measurement of same MDCK cell samples by QUEEN-37C and luciferase. Source data for Fig. 2C.

Supplementary Dataset S2

Single cell ATP measurement results of various cell types. Source data for Table 1.

Supplementary Dataset S3

Single cell ATP measurement of MDCK cells at different cell density. Soure data for Fig. 4E, F.

