## Supplemental Fig S1-S4, Supplemental Table S1-S4 for "Live cell imaging of metabolic heterogeneity by quantitative fluorescent ATP indicator protein, QUEEN-37C"

### Slide 1
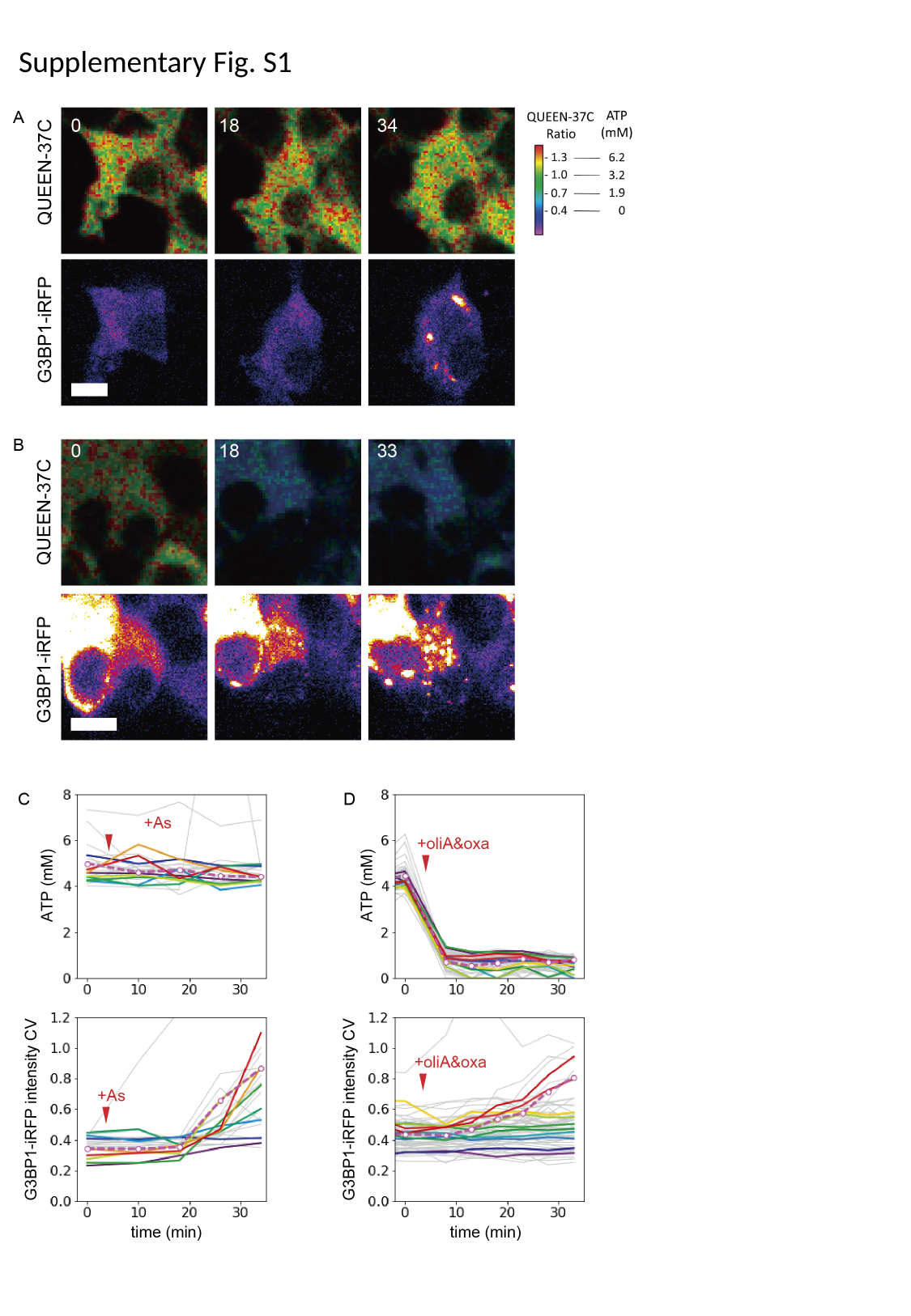

Supplementary Fig. S1

### Slide 2
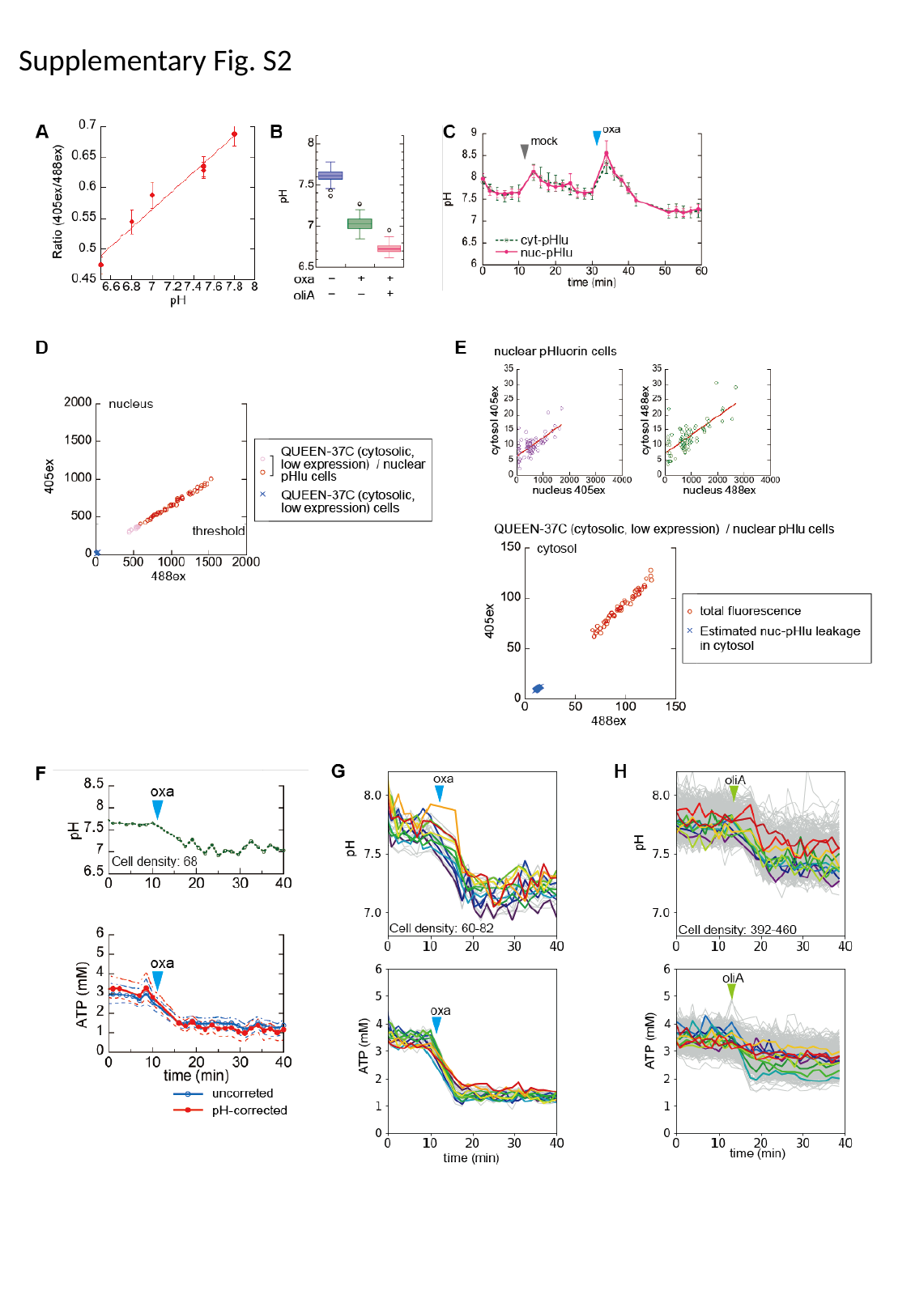

Supplementary Fig. S2

### Slide 3
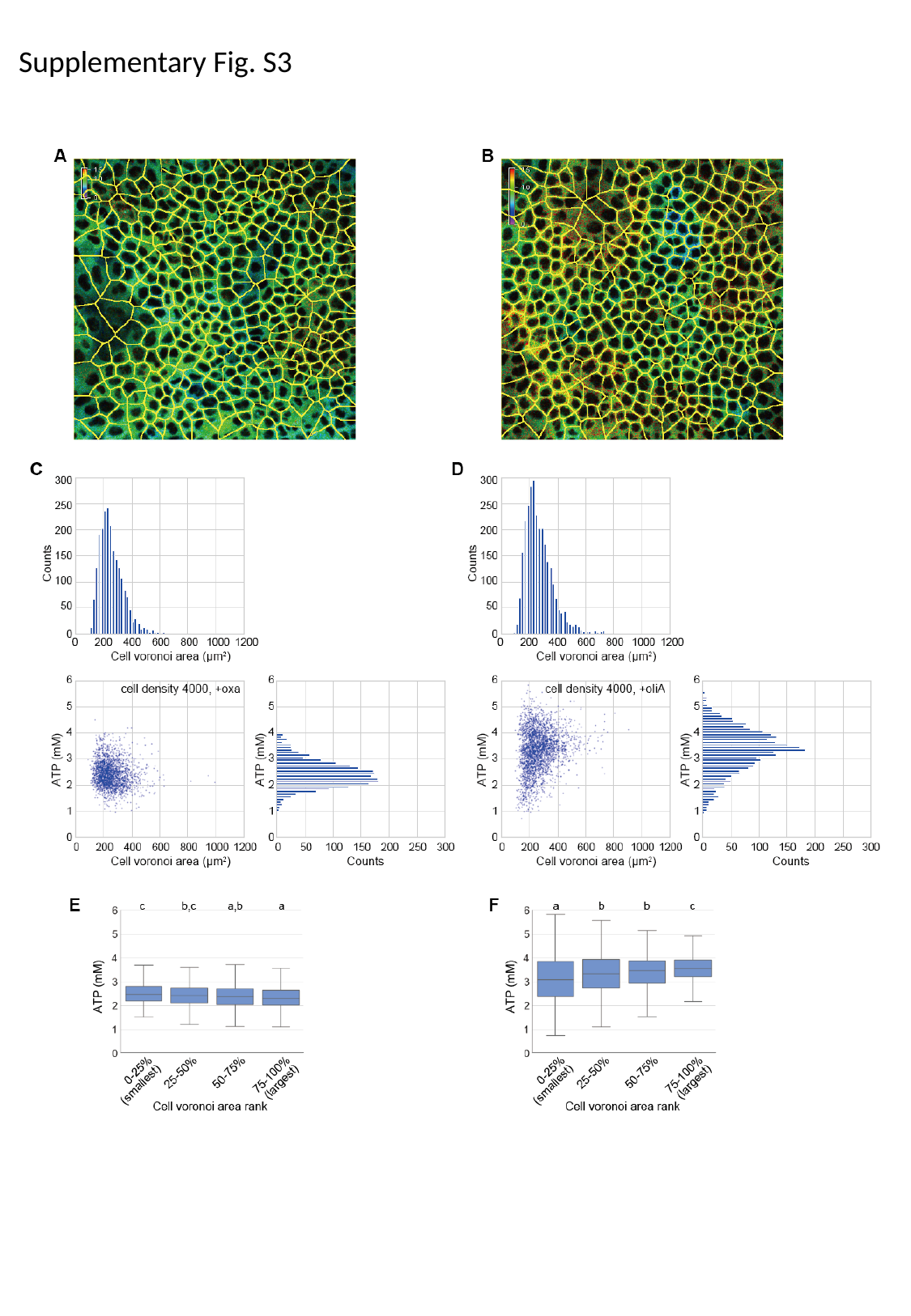

Supplementary Fig. S3

### Slide 4
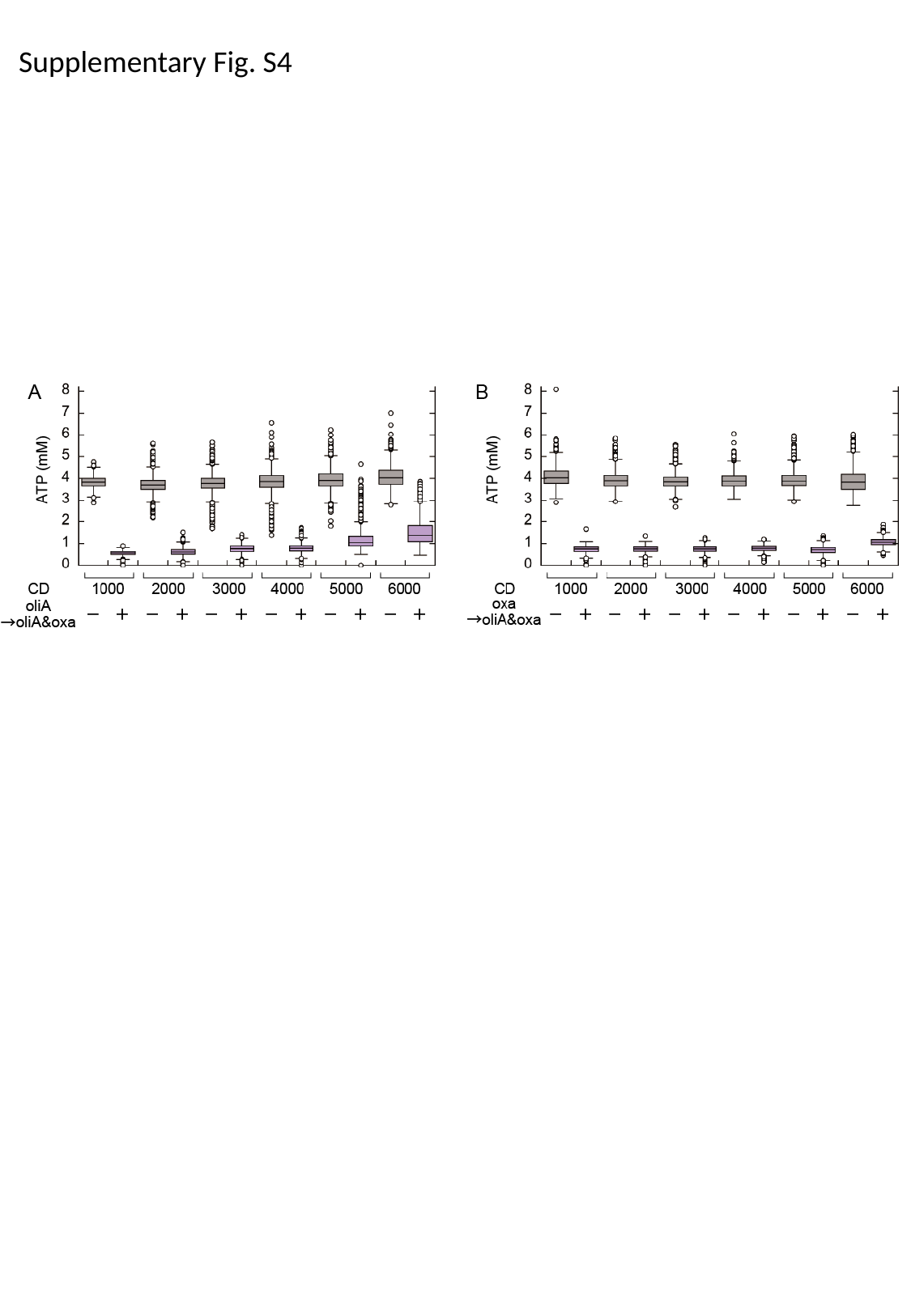

Supplementary Fig. S4

### Slide 5
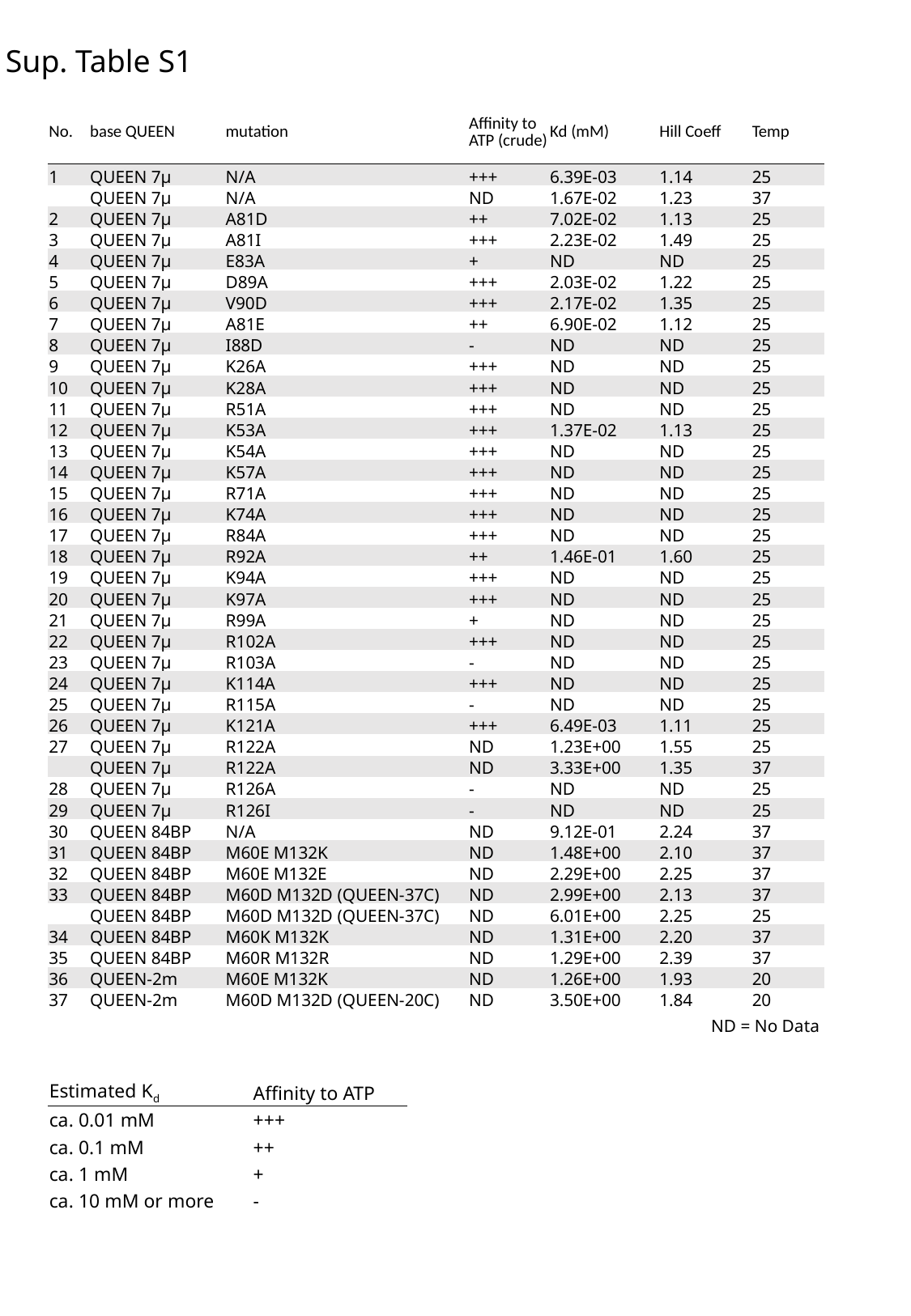

Sup. Table S1
| No. | base QUEEN | mutation | Affinity to ATP (crude) | Kd (mM) | Hill Coeff | Temp |
| --- | --- | --- | --- | --- | --- | --- |
| 1 | QUEEN 7µ | N/A | +++ | 6.39E-03 | 1.14 | 25 |
| | QUEEN 7µ | N/A | ND | 1.67E-02 | 1.23 | 37 |
| 2 | QUEEN 7µ | A81D | ++ | 7.02E-02 | 1.13 | 25 |
| 3 | QUEEN 7µ | A81I | +++ | 2.23E-02 | 1.49 | 25 |
| 4 | QUEEN 7µ | E83A | + | ND | ND | 25 |
| 5 | QUEEN 7µ | D89A | +++ | 2.03E-02 | 1.22 | 25 |
| 6 | QUEEN 7µ | V90D | +++ | 2.17E-02 | 1.35 | 25 |
| 7 | QUEEN 7µ | A81E | ++ | 6.90E-02 | 1.12 | 25 |
| 8 | QUEEN 7µ | I88D | - | ND | ND | 25 |
| 9 | QUEEN 7µ | K26A | +++ | ND | ND | 25 |
| 10 | QUEEN 7µ | K28A | +++ | ND | ND | 25 |
| 11 | QUEEN 7µ | R51A | +++ | ND | ND | 25 |
| 12 | QUEEN 7µ | K53A | +++ | 1.37E-02 | 1.13 | 25 |
| 13 | QUEEN 7µ | K54A | +++ | ND | ND | 25 |
| 14 | QUEEN 7µ | K57A | +++ | ND | ND | 25 |
| 15 | QUEEN 7µ | R71A | +++ | ND | ND | 25 |
| 16 | QUEEN 7µ | K74A | +++ | ND | ND | 25 |
| 17 | QUEEN 7µ | R84A | +++ | ND | ND | 25 |
| 18 | QUEEN 7µ | R92A | ++ | 1.46E-01 | 1.60 | 25 |
| 19 | QUEEN 7µ | K94A | +++ | ND | ND | 25 |
| 20 | QUEEN 7µ | K97A | +++ | ND | ND | 25 |
| 21 | QUEEN 7µ | R99A | + | ND | ND | 25 |
| 22 | QUEEN 7µ | R102A | +++ | ND | ND | 25 |
| 23 | QUEEN 7µ | R103A | - | ND | ND | 25 |
| 24 | QUEEN 7µ | K114A | +++ | ND | ND | 25 |
| 25 | QUEEN 7µ | R115A | - | ND | ND | 25 |
| 26 | QUEEN 7µ | K121A | +++ | 6.49E-03 | 1.11 | 25 |
| 27 | QUEEN 7µ | R122A | ND | 1.23E+00 | 1.55 | 25 |
| | QUEEN 7µ | R122A | ND | 3.33E+00 | 1.35 | 37 |
| 28 | QUEEN 7µ | R126A | - | ND | ND | 25 |
| 29 | QUEEN 7µ | R126I | - | ND | ND | 25 |
| 30 | QUEEN 84BP | N/A | ND | 9.12E-01 | 2.24 | 37 |
| 31 | QUEEN 84BP | M60E M132K | ND | 1.48E+00 | 2.10 | 37 |
| 32 | QUEEN 84BP | M60E M132E | ND | 2.29E+00 | 2.25 | 37 |
| 33 | QUEEN 84BP | M60D M132D (QUEEN-37C) | ND | 2.99E+00 | 2.13 | 37 |
| | QUEEN 84BP | M60D M132D (QUEEN-37C) | ND | 6.01E+00 | 2.25 | 25 |
| 34 | QUEEN 84BP | M60K M132K | ND | 1.31E+00 | 2.20 | 37 |
| 35 | QUEEN 84BP | M60R M132R | ND | 1.29E+00 | 2.39 | 37 |
| 36 | QUEEN-2m | M60E M132K | ND | 1.26E+00 | 1.93 | 20 |
| 37 | QUEEN-2m | M60D M132D (QUEEN-20C) | ND | 3.50E+00 | 1.84 | 20 |
ND = No Data
| Estimated K_d_ | Affinity to ATP |
| --- | --- |
| ca. 0.01 mM | +++ |
| ca. 0.1 mM | ++ |
| ca. 1 mM | + |
| ca. 10 mM or more | - |

### Slide 6
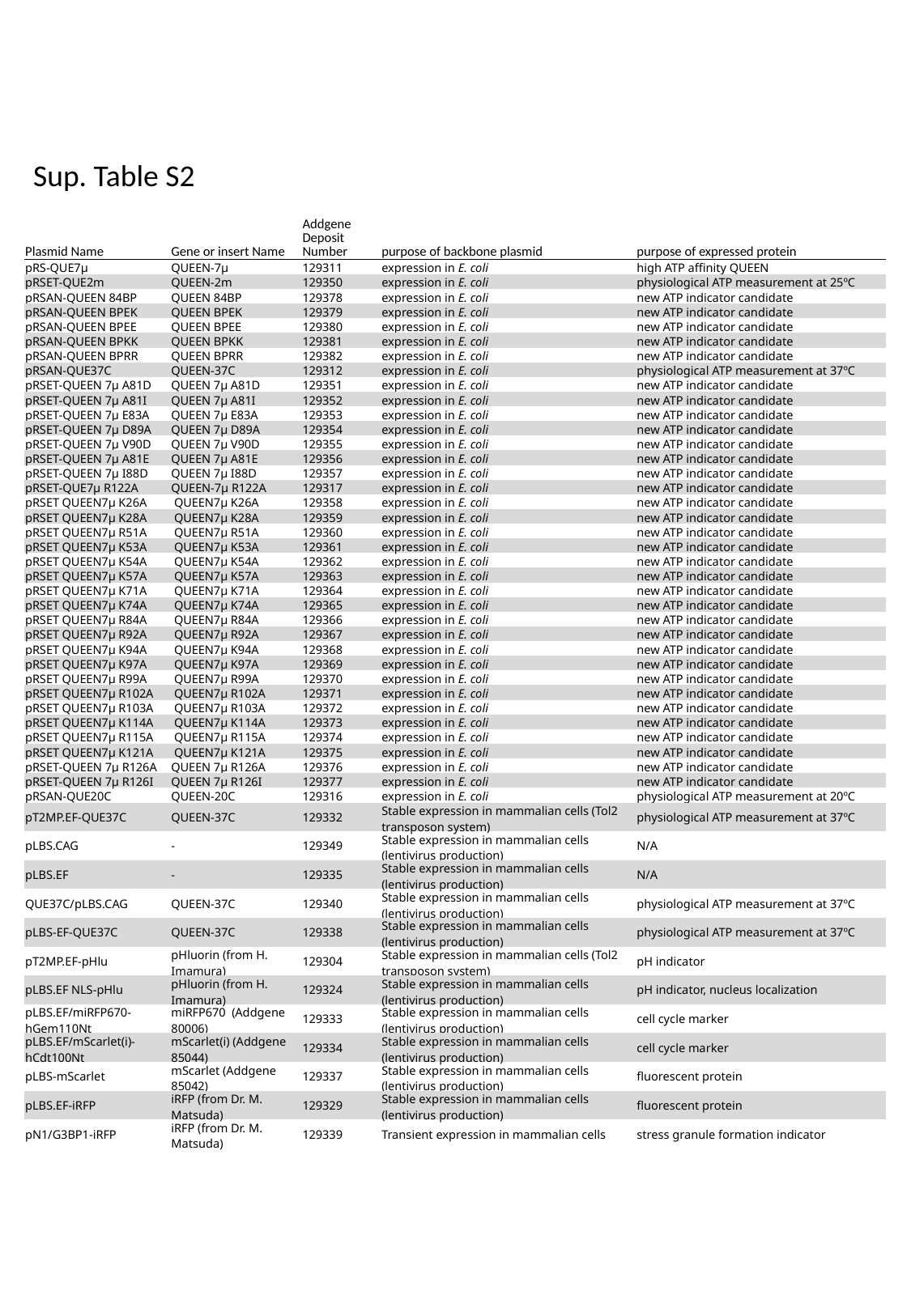

Sup. Table S2
| Plasmid Name | Gene or insert Name | Addgene Deposit Number | purpose of backbone plasmid | purpose of expressed protein |
| --- | --- | --- | --- | --- |
| pRS-QUE7µ | QUEEN-7µ | 129311 | expression in E. coli | high ATP affinity QUEEN |
| pRSET-QUE2m | QUEEN-2m | 129350 | expression in E. coli | physiological ATP measurement at 25ºC |
| pRSAN-QUEEN 84BP | QUEEN 84BP | 129378 | expression in E. coli | new ATP indicator candidate |
| pRSAN-QUEEN BPEK | QUEEN BPEK | 129379 | expression in E. coli | new ATP indicator candidate |
| pRSAN-QUEEN BPEE | QUEEN BPEE | 129380 | expression in E. coli | new ATP indicator candidate |
| pRSAN-QUEEN BPKK | QUEEN BPKK | 129381 | expression in E. coli | new ATP indicator candidate |
| pRSAN-QUEEN BPRR | QUEEN BPRR | 129382 | expression in E. coli | new ATP indicator candidate |
| pRSAN-QUE37C | QUEEN-37C | 129312 | expression in E. coli | physiological ATP measurement at 37ºC |
| pRSET-QUEEN 7µ A81D | QUEEN 7µ A81D | 129351 | expression in E. coli | new ATP indicator candidate |
| pRSET-QUEEN 7µ A81I | QUEEN 7µ A81I | 129352 | expression in E. coli | new ATP indicator candidate |
| pRSET-QUEEN 7µ E83A | QUEEN 7µ E83A | 129353 | expression in E. coli | new ATP indicator candidate |
| pRSET-QUEEN 7µ D89A | QUEEN 7µ D89A | 129354 | expression in E. coli | new ATP indicator candidate |
| pRSET-QUEEN 7µ V90D | QUEEN 7µ V90D | 129355 | expression in E. coli | new ATP indicator candidate |
| pRSET-QUEEN 7µ A81E | QUEEN 7µ A81E | 129356 | expression in E. coli | new ATP indicator candidate |
| pRSET-QUEEN 7µ I88D | QUEEN 7µ I88D | 129357 | expression in E. coli | new ATP indicator candidate |
| pRSET-QUE7µ R122A | QUEEN-7µ R122A | 129317 | expression in E. coli | new ATP indicator candidate |
| pRSET QUEEN7µ K26A | QUEEN7µ K26A | 129358 | expression in E. coli | new ATP indicator candidate |
| pRSET QUEEN7µ K28A | QUEEN7µ K28A | 129359 | expression in E. coli | new ATP indicator candidate |
| pRSET QUEEN7µ R51A | QUEEN7µ R51A | 129360 | expression in E. coli | new ATP indicator candidate |
| pRSET QUEEN7µ K53A | QUEEN7µ K53A | 129361 | expression in E. coli | new ATP indicator candidate |
| pRSET QUEEN7µ K54A | QUEEN7µ K54A | 129362 | expression in E. coli | new ATP indicator candidate |
| pRSET QUEEN7µ K57A | QUEEN7µ K57A | 129363 | expression in E. coli | new ATP indicator candidate |
| pRSET QUEEN7µ K71A | QUEEN7µ K71A | 129364 | expression in E. coli | new ATP indicator candidate |
| pRSET QUEEN7µ K74A | QUEEN7µ K74A | 129365 | expression in E. coli | new ATP indicator candidate |
| pRSET QUEEN7µ R84A | QUEEN7µ R84A | 129366 | expression in E. coli | new ATP indicator candidate |
| pRSET QUEEN7µ R92A | QUEEN7µ R92A | 129367 | expression in E. coli | new ATP indicator candidate |
| pRSET QUEEN7µ K94A | QUEEN7µ K94A | 129368 | expression in E. coli | new ATP indicator candidate |
| pRSET QUEEN7µ K97A | QUEEN7µ K97A | 129369 | expression in E. coli | new ATP indicator candidate |
| pRSET QUEEN7µ R99A | QUEEN7µ R99A | 129370 | expression in E. coli | new ATP indicator candidate |
| pRSET QUEEN7µ R102A | QUEEN7µ R102A | 129371 | expression in E. coli | new ATP indicator candidate |
| pRSET QUEEN7µ R103A | QUEEN7µ R103A | 129372 | expression in E. coli | new ATP indicator candidate |
| pRSET QUEEN7µ K114A | QUEEN7µ K114A | 129373 | expression in E. coli | new ATP indicator candidate |
| pRSET QUEEN7µ R115A | QUEEN7µ R115A | 129374 | expression in E. coli | new ATP indicator candidate |
| pRSET QUEEN7µ K121A | QUEEN7µ K121A | 129375 | expression in E. coli | new ATP indicator candidate |
| pRSET-QUEEN 7µ R126A | QUEEN 7µ R126A | 129376 | expression in E. coli | new ATP indicator candidate |
| pRSET-QUEEN 7µ R126I | QUEEN 7µ R126I | 129377 | expression in E. coli | new ATP indicator candidate |
| pRSAN-QUE20C | QUEEN-20C | 129316 | expression in E. coli | physiological ATP measurement at 20ºC |
| pT2MP.EF-QUE37C | QUEEN-37C | 129332 | Stable expression in mammalian cells (Tol2 transposon system) | physiological ATP measurement at 37ºC |
| pLBS.CAG | - | 129349 | Stable expression in mammalian cells (lentivirus production) | N/A |
| pLBS.EF | - | 129335 | Stable expression in mammalian cells (lentivirus production) | N/A |
| QUE37C/pLBS.CAG | QUEEN-37C | 129340 | Stable expression in mammalian cells (lentivirus production) | physiological ATP measurement at 37ºC |
| pLBS-EF-QUE37C | QUEEN-37C | 129338 | Stable expression in mammalian cells (lentivirus production) | physiological ATP measurement at 37ºC |
| pT2MP.EF-pHlu | pHluorin (from H. Imamura) | 129304 | Stable expression in mammalian cells (Tol2 transposon system) | pH indicator |
| pLBS.EF NLS-pHlu | pHluorin (from H. Imamura) | 129324 | Stable expression in mammalian cells (lentivirus production) | pH indicator, nucleus localization |
| pLBS.EF/miRFP670-hGem110Nt | miRFP670 (Addgene 80006) | 129333 | Stable expression in mammalian cells (lentivirus production) | cell cycle marker |
| pLBS.EF/mScarlet(i)-hCdt100Nt | mScarlet(i) (Addgene 85044) | 129334 | Stable expression in mammalian cells (lentivirus production) | cell cycle marker |
| pLBS-mScarlet | mScarlet (Addgene 85042) | 129337 | Stable expression in mammalian cells (lentivirus production) | fluorescent protein |
| pLBS.EF-iRFP | iRFP (from Dr. M. Matsuda) | 129329 | Stable expression in mammalian cells (lentivirus production) | fluorescent protein |
| pN1/G3BP1-iRFP | iRFP (from Dr. M. Matsuda) | 129339 | Transient expression in mammalian cells | stress granule formation indicator |

### Slide 7
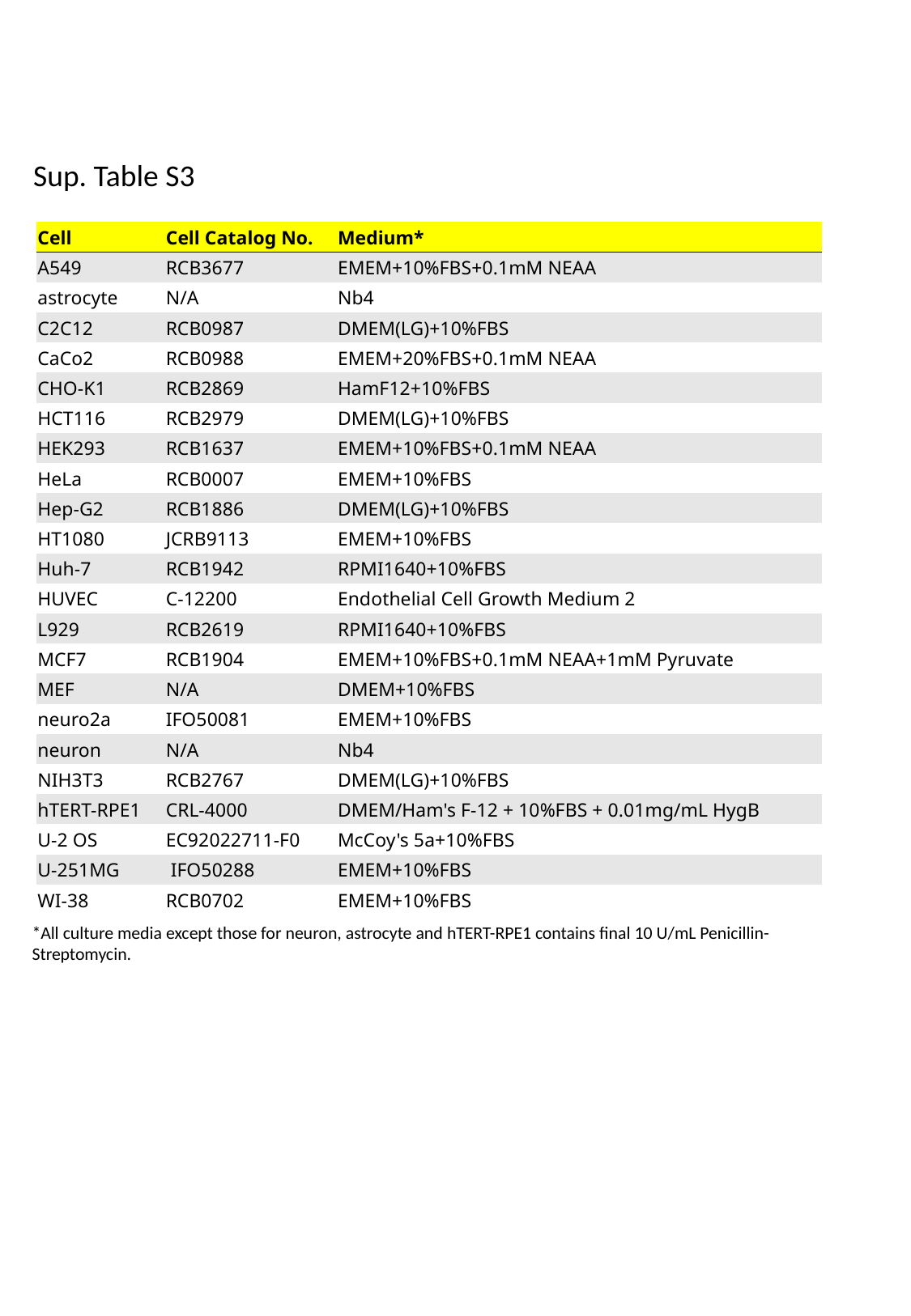

Sup. Table S3
| Cell | Cell Catalog No. | Medium\* |
| --- | --- | --- |
| A549 | RCB3677 | EMEM+10%FBS+0.1mM NEAA |
| astrocyte | N/A | Nb4 |
| C2C12 | RCB0987 | DMEM(LG)+10%FBS |
| CaCo2 | RCB0988 | EMEM+20%FBS+0.1mM NEAA |
| CHO-K1 | RCB2869 | HamF12+10%FBS |
| HCT116 | RCB2979 | DMEM(LG)+10%FBS |
| HEK293 | RCB1637 | EMEM+10%FBS+0.1mM NEAA |
| HeLa | RCB0007 | EMEM+10%FBS |
| Hep-G2 | RCB1886 | DMEM(LG)+10%FBS |
| HT1080 | JCRB9113 | EMEM+10%FBS |
| Huh-7 | RCB1942 | RPMI1640+10%FBS |
| HUVEC | C-12200 | Endothelial Cell Growth Medium 2 |
| L929 | RCB2619 | RPMI1640+10%FBS |
| MCF7 | RCB1904 | EMEM+10%FBS+0.1mM NEAA+1mM Pyruvate |
| MEF | N/A | DMEM+10%FBS |
| neuro2a | IFO50081 | EMEM+10%FBS |
| neuron | N/A | Nb4 |
| NIH3T3 | RCB2767 | DMEM(LG)+10%FBS |
| hTERT-RPE1 | CRL-4000 | DMEM/Ham's F-12 + 10%FBS + 0.01mg/mL HygB |
| U-2 OS | EC92022711-F0 | McCoy's 5a+10%FBS |
| U-251MG | IFO50288 | EMEM+10%FBS |
| WI-38 | RCB0702 | EMEM+10%FBS |
*All culture media except those for neuron, astrocyte and hTERT-RPE1 contains final 10 U/mL Penicillin-Streptomycin.

### Slide 8
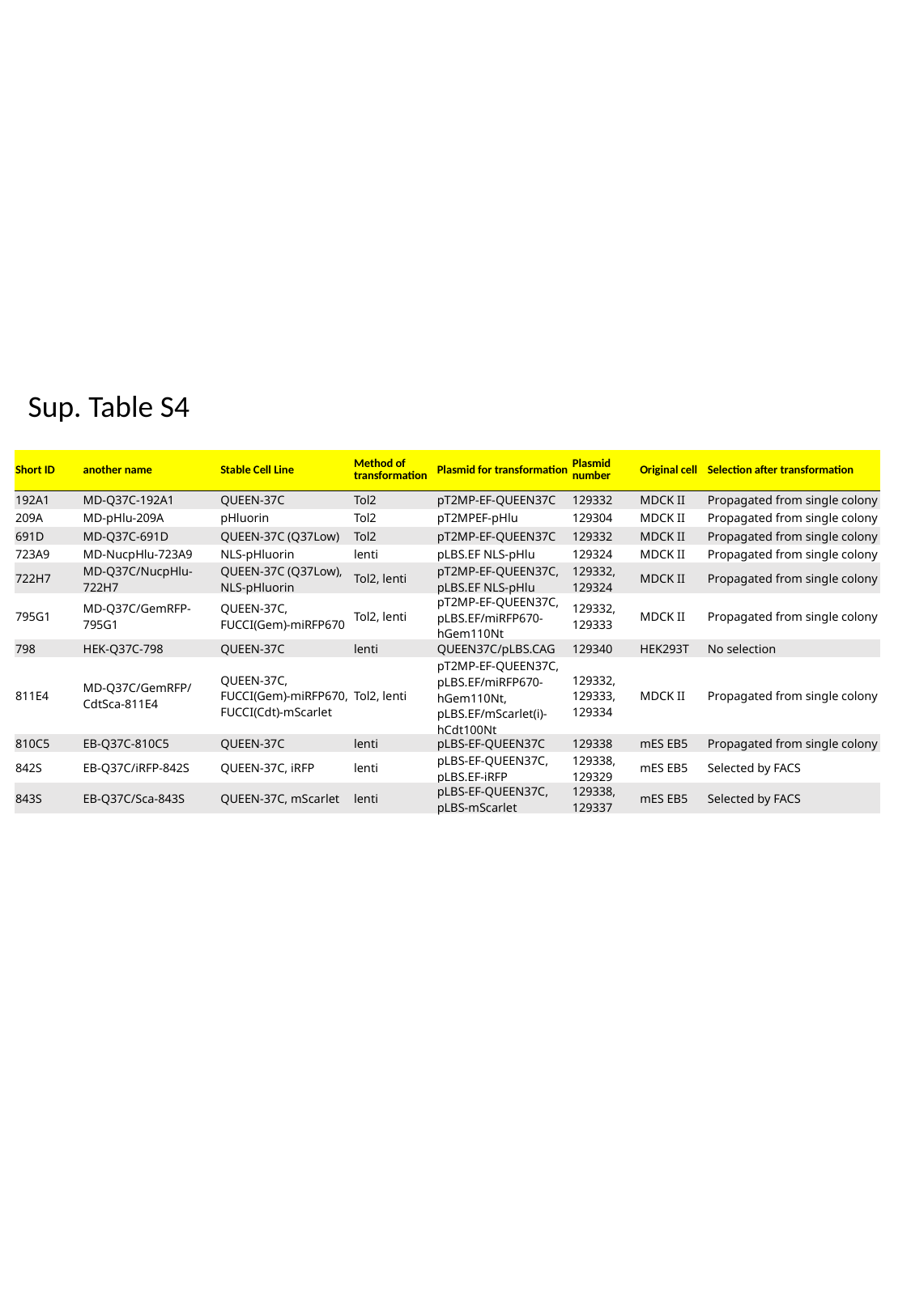

Sup. Table S4
| Short ID | another name | Stable Cell Line | Method of transformation | Plasmid for transformation | Plasmid number | Original cell | Selection after transformation |
| --- | --- | --- | --- | --- | --- | --- | --- |
| 192A1 | MD-Q37C-192A1 | QUEEN-37C | Tol2 | pT2MP-EF-QUEEN37C | 129332 | MDCK II | Propagated from single colony |
| 209A | MD-pHlu-209A | pHluorin | Tol2 | pT2MPEF-pHlu | 129304 | MDCK II | Propagated from single colony |
| 691D | MD-Q37C-691D | QUEEN-37C (Q37Low) | Tol2 | pT2MP-EF-QUEEN37C | 129332 | MDCK II | Propagated from single colony |
| 723A9 | MD-NucpHlu-723A9 | NLS-pHluorin | lenti | pLBS.EF NLS-pHlu | 129324 | MDCK II | Propagated from single colony |
| 722H7 | MD-Q37C/NucpHlu-722H7 | QUEEN-37C (Q37Low), NLS-pHluorin | Tol2, lenti | pT2MP-EF-QUEEN37C, pLBS.EF NLS-pHlu | 129332, 129324 | MDCK II | Propagated from single colony |
| 795G1 | MD-Q37C/GemRFP-795G1 | QUEEN-37C, FUCCI(Gem)-miRFP670 | Tol2, lenti | pT2MP-EF-QUEEN37C, pLBS.EF/miRFP670-hGem110Nt | 129332, 129333 | MDCK II | Propagated from single colony |
| 798 | HEK-Q37C-798 | QUEEN-37C | lenti | QUEEN37C/pLBS.CAG | 129340 | HEK293T | No selection |
| 811E4 | MD-Q37C/GemRFP/CdtSca-811E4 | QUEEN-37C, FUCCI(Gem)-miRFP670, FUCCI(Cdt)-mScarlet | Tol2, lenti | pT2MP-EF-QUEEN37C, pLBS.EF/miRFP670-hGem110Nt, pLBS.EF/mScarlet(i)-hCdt100Nt | 129332, 129333, 129334 | MDCK II | Propagated from single colony |
| 810C5 | EB-Q37C-810C5 | QUEEN-37C | lenti | pLBS-EF-QUEEN37C | 129338 | mES EB5 | Propagated from single colony |
| 842S | EB-Q37C/iRFP-842S | QUEEN-37C, iRFP | lenti | pLBS-EF-QUEEN37C, pLBS.EF-iRFP | 129338, 129329 | mES EB5 | Selected by FACS |
| 843S | EB-Q37C/Sca-843S | QUEEN-37C, mScarlet | lenti | pLBS-EF-QUEEN37C, pLBS-mScarlet | 129338, 129337 | mES EB5 | Selected by FACS |
